# Investigating and Identifying Traditional Medicinal Plant Antioxidants– New Frontiers in Treating Age-related Maladies Associated with Oxidative Stress

**DOI:** 10.1101/2023.11.06.565860

**Authors:** Finan Gammell, Dean Gangji, Elizabeth Leibovitz, Matthew J Bertin

**Affiliations:** East Greenwich High School; Department of Biomedical and Pharmaceutical Sciences, College of Pharmacy, University of Rhode Island

## Abstract

Oxidative stress is regarded as an imbalance between pro-oxidant and antioxidant species that can result in cellular inflammation associated with illnesses including cancers and cognitive decline. This study investigated medicinal plant extracts for the presence of natural bioactive compounds to explore their antioxidant properties and potential therapeutic relevance in conditions related to oxidative stress. Extracts were created using water as a solvent and compared in triplicate to the positive control quercetin (1 mg/mL) in a 2,2-diphenyl-1-picrylhydrazyl (DPPH) assay; all extracts were serially diluted from 10 mg/mL creating 8 distinct concentrations. Absorbance readings were compared before and after DPPH administration. At the highest concentration, all extracts demonstrated absorbance change factors <10% relative to the DPPH negative control. This result clearly demonstrated antioxidant activity, as lesser relative absorbance changes indicated greater neutralization of violet DPPH into lighter diphenylpicryl hydrazine via free radical scavenging. Most notably, at lowest concentration, *Averrhoa carambola* and *Psidium guajava* represented the least relative absorbance changes at 14.17% and 10.39%, rivaling quercetin’s antioxidant activity– a statistically significant result (p < 0.05). LC-MS/MS analysis of these most potent extracts identified quercetin and kaempferol in *Psidium guajava* and vitexin-2’’-O-rhamnoside, isovitexin, saponarin, and an additional flavonoid glycoside in *Averrhoa carambola*. Such compounds may offer potential pharmacological effects, for example, vitexin-2’’-O-rhamnoside and isovitexin have demonstrated anticancer, antidiabetic, neuroprotective, and cardioprotective properties in part through oxidative stress reduction. Future research should corroborate compound effects in human cell culture models, as prior studies are often limited to non-human tissues.

**INTRODUCTION:** A medicinal plant is a plant that “contains substances that can be used for therapeutic purposes or which are precursors for the synthesis of useful drugs.”^1^ Medicinal plants contain different classes of potentially bioactive compounds, including, but not limited to peptides, alkaloids, phenylpropanoids, polyketides, and terpenoids.^2^ Such compounds have alleged and in certain cases documented anticancer, anti-inflammatory, antioxidant, antibacterial, antipyretic, vasodilatory, stimulant, and/or more properties that have been historically used in traditional medicine.^1,3^ Evidence suggests that medicinal plants have been used for at least 60,000 years in places including what would be modern-day Iraq.^4^ Medicinal plants are still used all around the world today; Dr. Tedros Adhanom Ghebreyesus of the WHO says “For many millions of people around the world, traditional medicine is the first port of call to treat many diseases.”^5^ In addition, the prevalence of medicinal plants is evident in developed countries too, with the medicinal plant trade between India and China accounting for two to five billion dollars annually.^1^

However, many of these traditional treatments receive significant scrutiny from modern medicine and society’s decreasing reliance on plant-derived drugs. While such scrutiny proves accurate in many cases, several traditional medicinal plants have been shown to offer bioactive compounds that are used as a standard medical practice. This includes morphine, “a natural alkaloid that is derived from resin extracts from the seeds of the opium poppy,”^3^ which is utilized in clinical settings for its profound analgesic effects,^3^ and quinine, an antimalarial drug derived from *Cinchona officinalis* bark.^6^ The current research project presented herein will serve to test traditional medicinal plant compounds to determine if their claims of relevance can be substantiated and to explore their potential effects on biological organisms, specifically regarding their antioxidant properties and ability to eliminate free radicals. Free radicals are unstable molecular species with unpaired electrons,^7^ and can contribute to oxidative stress in organisms. Oxidative stress “is considered as an imbalance between pro- and antioxidant species, which results in molecular and cellular damage.”^8^ Examples of known antioxidants include the polyphenol quercetin, which is often used as a positive control for its relative strength in reducing free radicals.^9^ Demonstrated efficacy of these medicinal plants could provide potential preventative therapies for age-related illnesses and other maladies associated with oxidative stress, including cancers and cognitive decline.^8,9^

*RESEARCH QUESTIONS AND PURPOSE:* Do the traditional medicinal plants under investigation contain bioactive compounds with antioxidant properties capable of neutralizing free radicals in a 2,2-diphenyl-1-picrylhydrazyl (DPPH) assay? How does the efficacy of the plant extracts compare to the positive control, quercetin, and the associated samples in terms of antioxidant activity as measured by the DPPH assay? Do the extract compounds identified via LC-MS/MS analysis offer potential therapeutic relevance in modern medicine? With our research, we aimed to identify bioactive compounds within medicinal plants, determine if such medicinal plants and their bioactive compounds have applications in modern medical treatments, and ensure medicinal plants are being used safely and accurately under the support of sound scientific evidence.

## MATERIALS AND METHODS

Aerial Specimens (Leaves, twigs, and branches) collected from:

-*Camellia sinensis*
-*Averrhoa carambola*
-*Salvia rosmarinus*
-*Laurus nobilis*
-*Psidium guajava*

*Other materials:*

-Pure MilleQ water
-250 mL Erlenmeyer flasks
-Foil
-Graduated cylinder
-Weigh boats
-Balance
-Scissors & forceps
-Ethanol
-2,2-diphenyl-1-picrylhydrazyl (DPPH)
-96 well clear bottom plates
-Solvent reservoirs
-Multi-channel pipette
-p100 pipette tips
-Spectromax plate reader
-Quercetin (positive control in antioxidant assays)
-HPLC vials
-C18 SPE cartridges
-ThermoScientific LTQ XL mass spectrometer with Dionex Ultimate 3000 HPLC system

### Part I: Make Extract/Tea

Plant material was harvested from the URI Heber W. Youngken Jr. Medicinal Garden Greenhouse. Specimens consisted of aerial parts; leaves, twigs, and small branches. In the laboratory, the plant material was dissected and 5.00g portions (4.00g leaves; 1.00g twigs/small branches) of plant specimens were weighed and added to an Erlenmeyer flask. Next, 50 mL of water was poured onto each sample of plant material, and the flask was sealed with foil. Extractions took place under the chemical fume hood for 6 days after which a 15 mL aliquot of each sample was removed, transferred to a pre-weighed vial, and then dried via evaporation with inert nitrogen to determine the exact dried extract weight (Figure 1).

**Figure 1.**
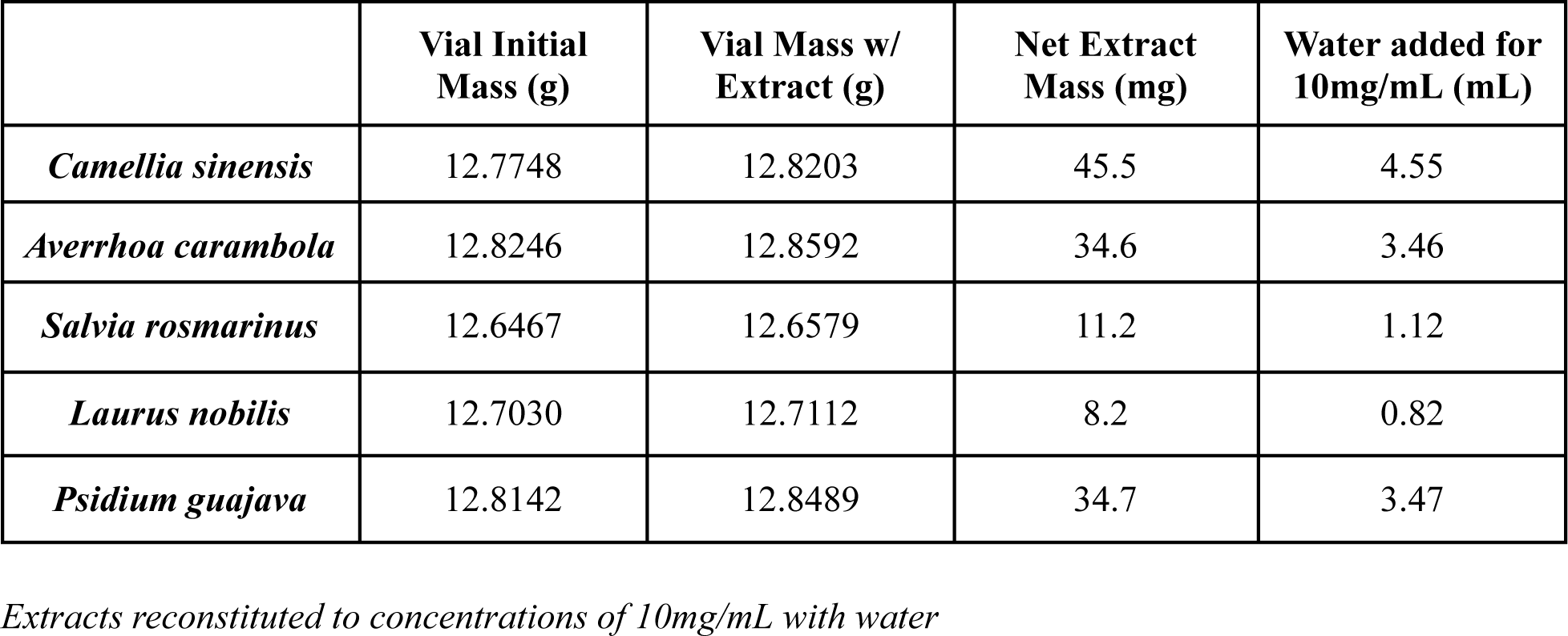
Extract Masses and Reconstitution.

### Part II: Test Extracts through DPPH assay

Each water extract was weighed and reconstituted to a 10 mg/mL concentration using water (Figure 1). A positive control mixture of quercetin was prepared in ethanol at 1 mg/mL and a solution of 200 µM of DPPH was also made in ethanol. To a 96-well plate, concentrations of quercetin (1 mg/mL) and the plant extracts (10 mg/mL) were added to the top row of the assay plate. Thus, each concentration of positive control and test sample was tested in triplicate. Next, using a multichannel pipette or a standard pipette, we aliquoted 100 µl of water into each well of rows B-H (Columns 3-11) of a 96-well plate. Next, we serially diluted the positive control and test extract samples by taking 100 µl of the control and extracts from row A and pipetting them into the 100 µl of water in row B. We mixed them well by pipetting up and down several times, then transferred 100 µl from row B into row C and repeated until row H had been thoroughly mixed, then discarded the remaining 100 µl from row H wells into the first reservoir that was used for the water (See Figures 2.a, 2.b, and 2.c, for plate maps). Following the serial dilution, we returned to the bench and poured the pre-diluted DPPH solution into the second solvent reservoir, and we pipetted 100 µl of DPPH into each well. We covered the plate in foil and waited for 30 min for the assay to resolve (See Figures 3.a, 3.b, and 3.c). Finally, we analyzed the plate in a Spectromax plate reader measuring the absorbance at 517 nm. Bar graphs were made with a concentration on the x-axis and absorbance on the y-axis.

**Figure 2.a.**
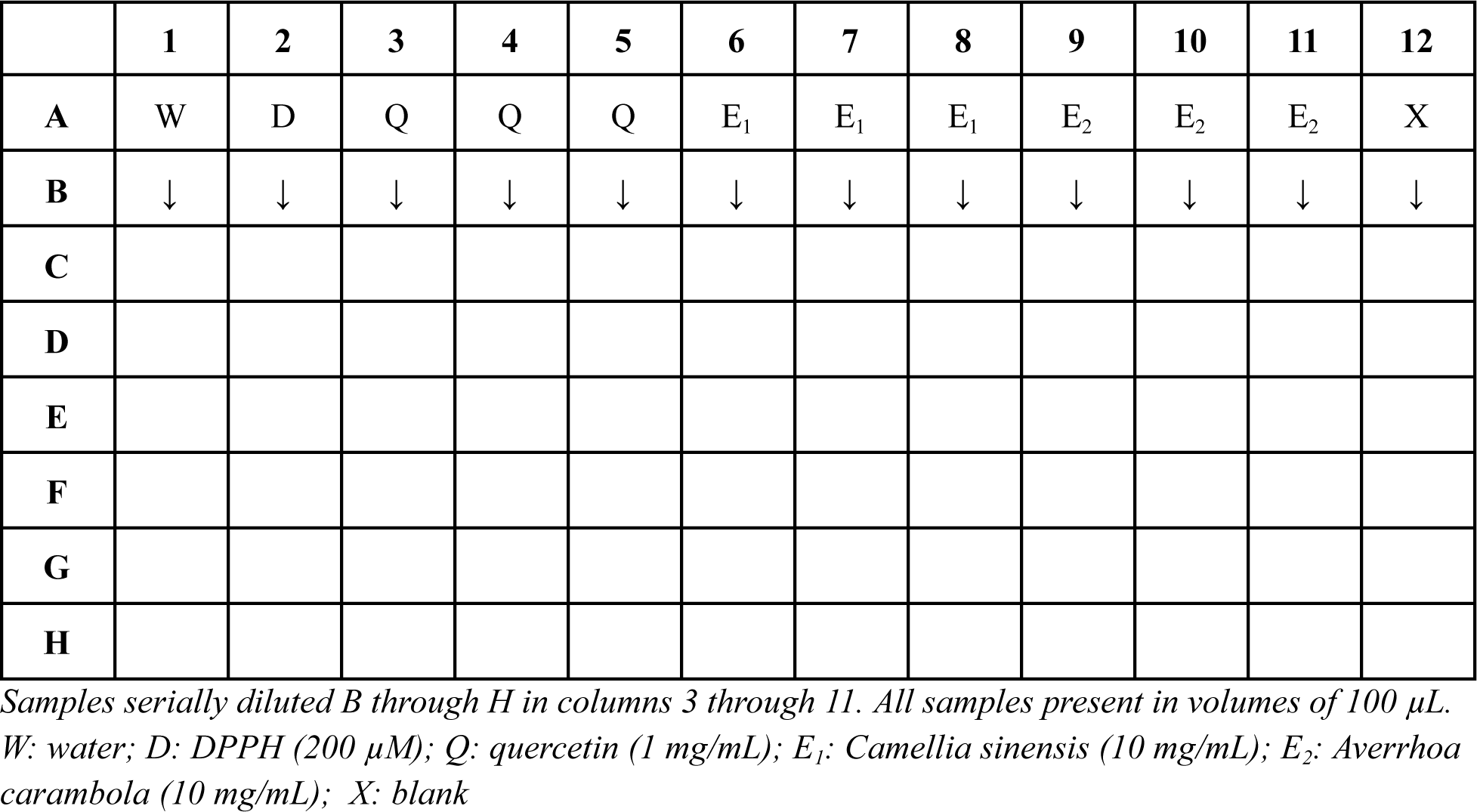
DPPH Assay Plate Map A (*Camellia sinensis* & *Averrhoa carambola*)

**Figure 2.b.**
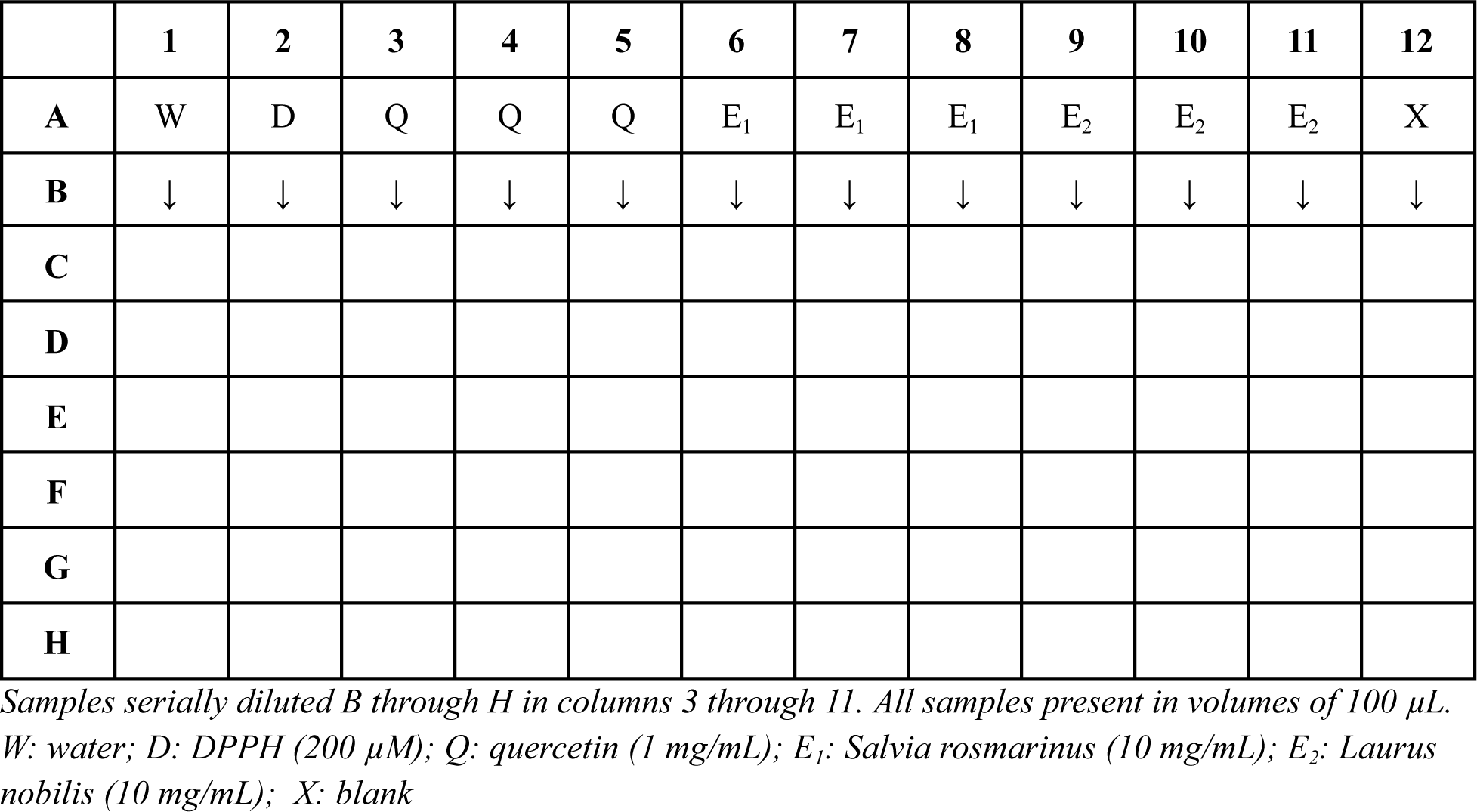
DPPH Assay Plate Map B (*Salvia rosmarinus* & *Laurus nobilis*)

**Figure 2.c.**
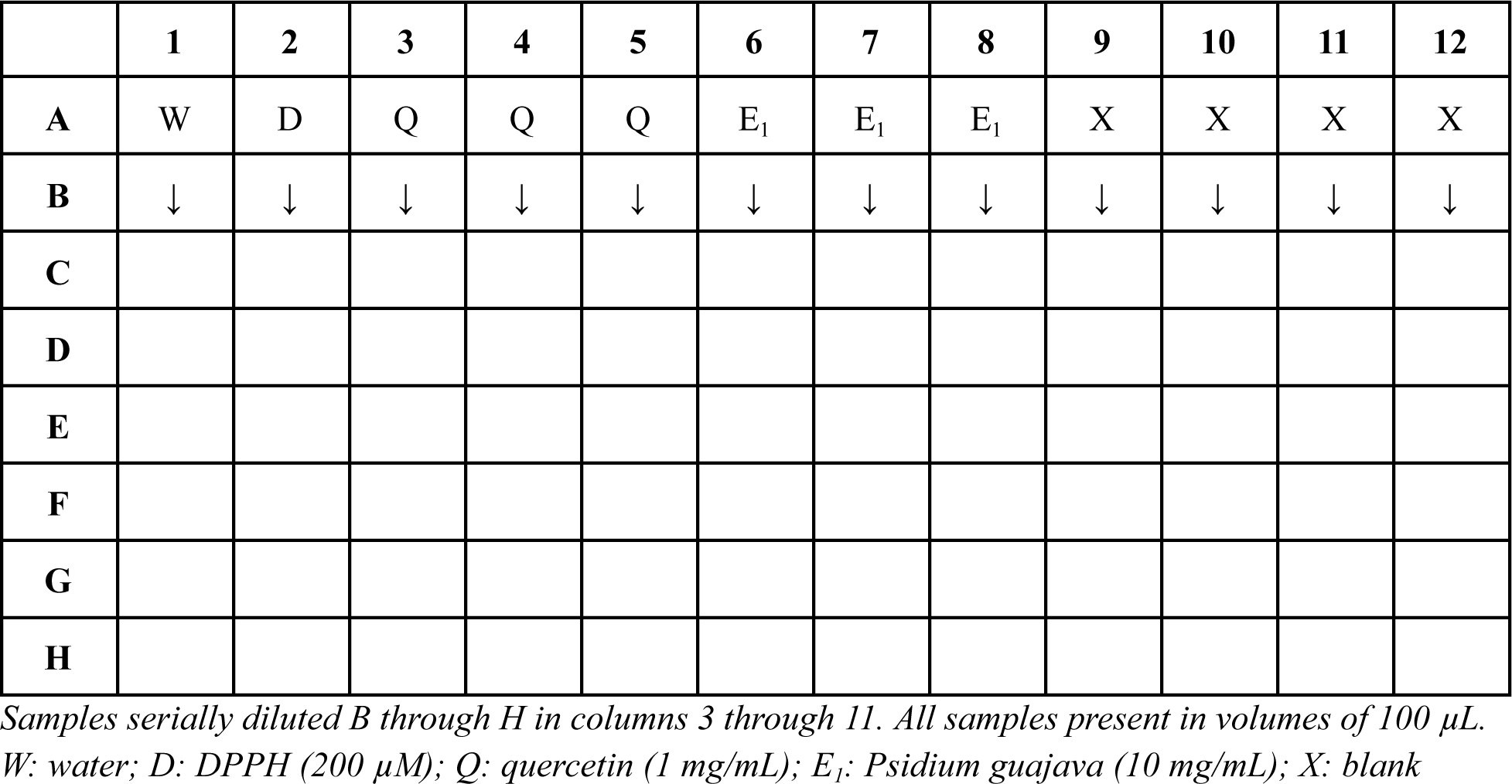
DPPH Assay Plate Map C (*Psidium guajava*)

**Figure 3.a.**
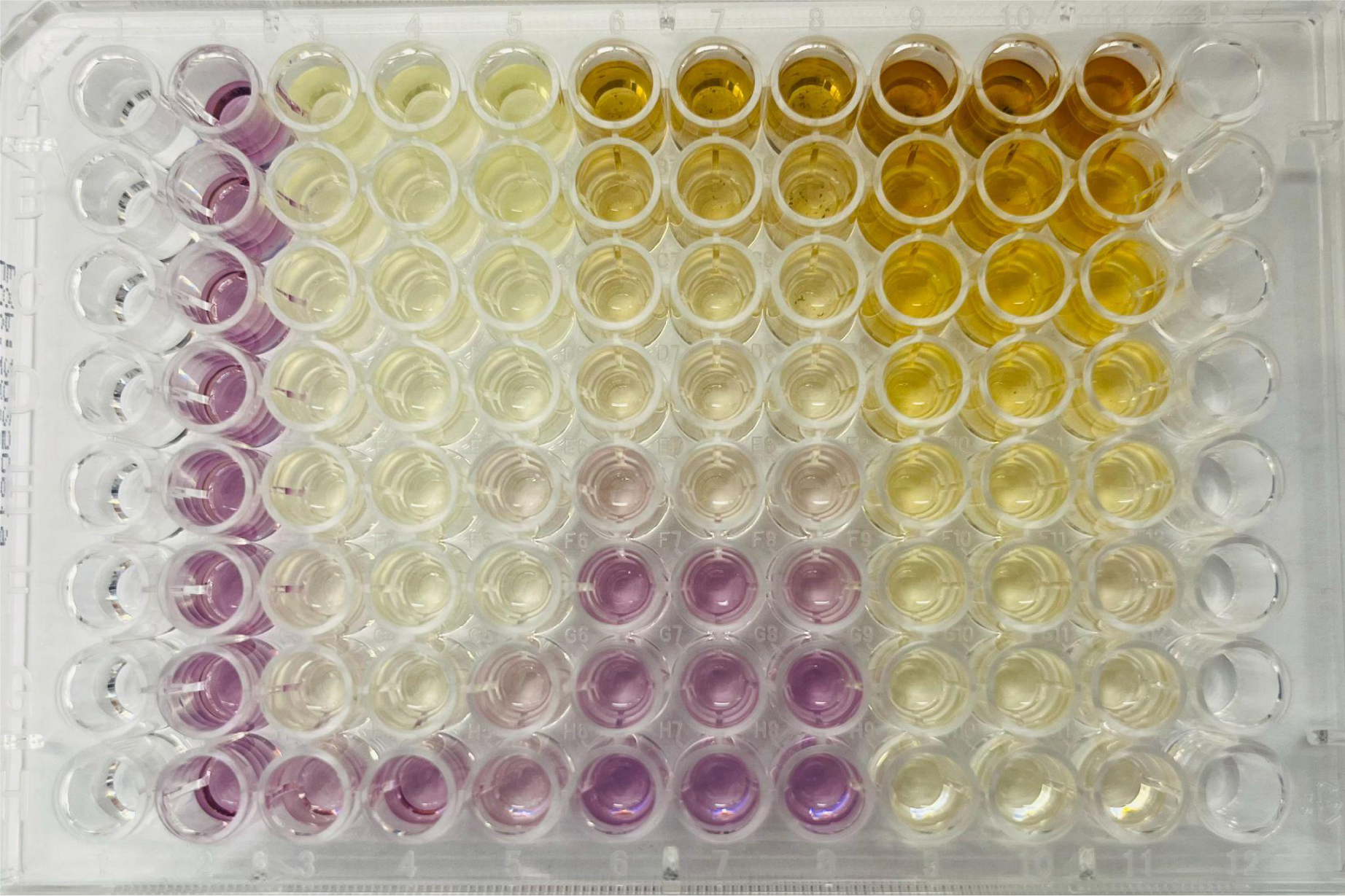
DPPH Assay A.

**Figure 3.b.**
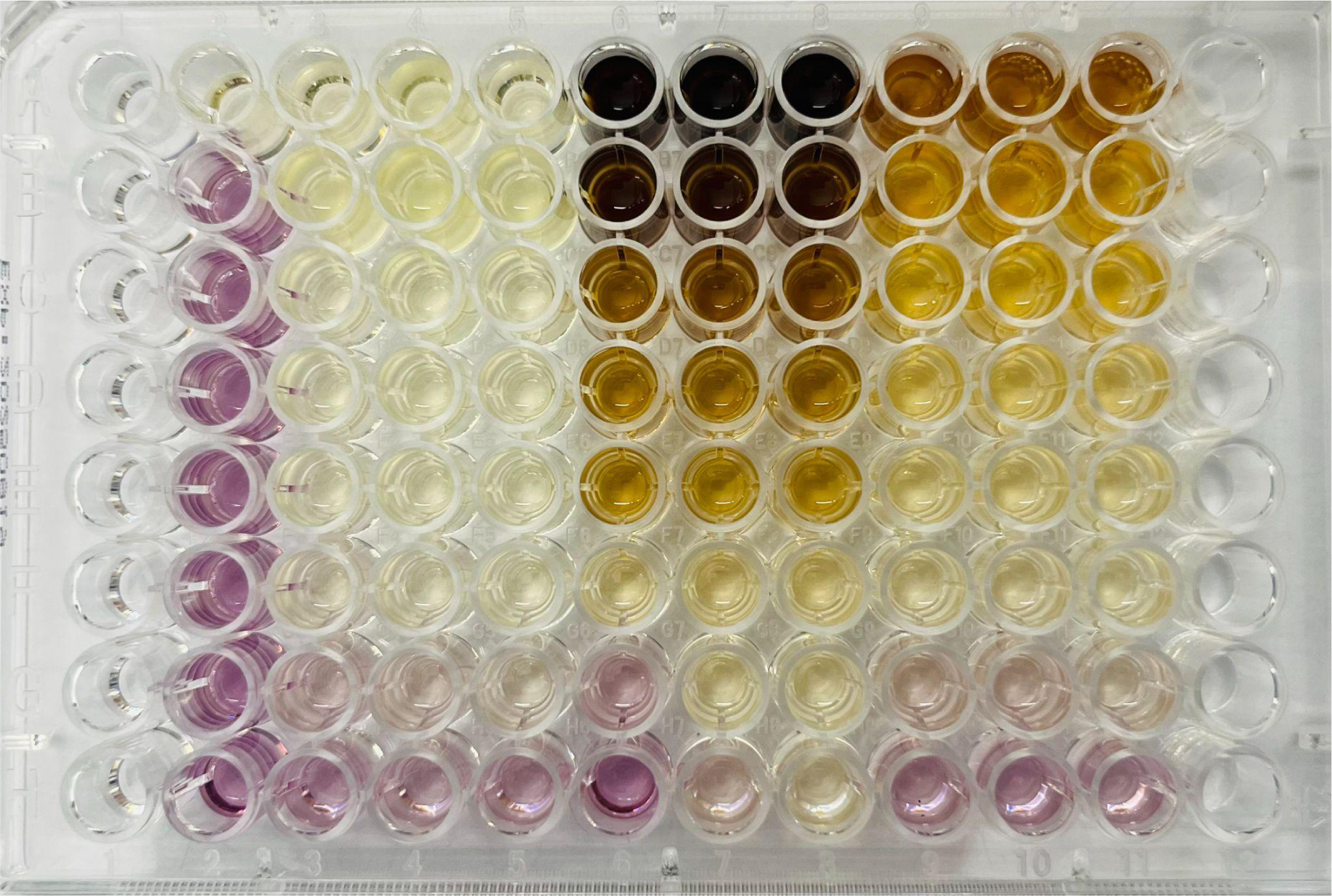
DPPH Assay B.

**Figure 3.c.**
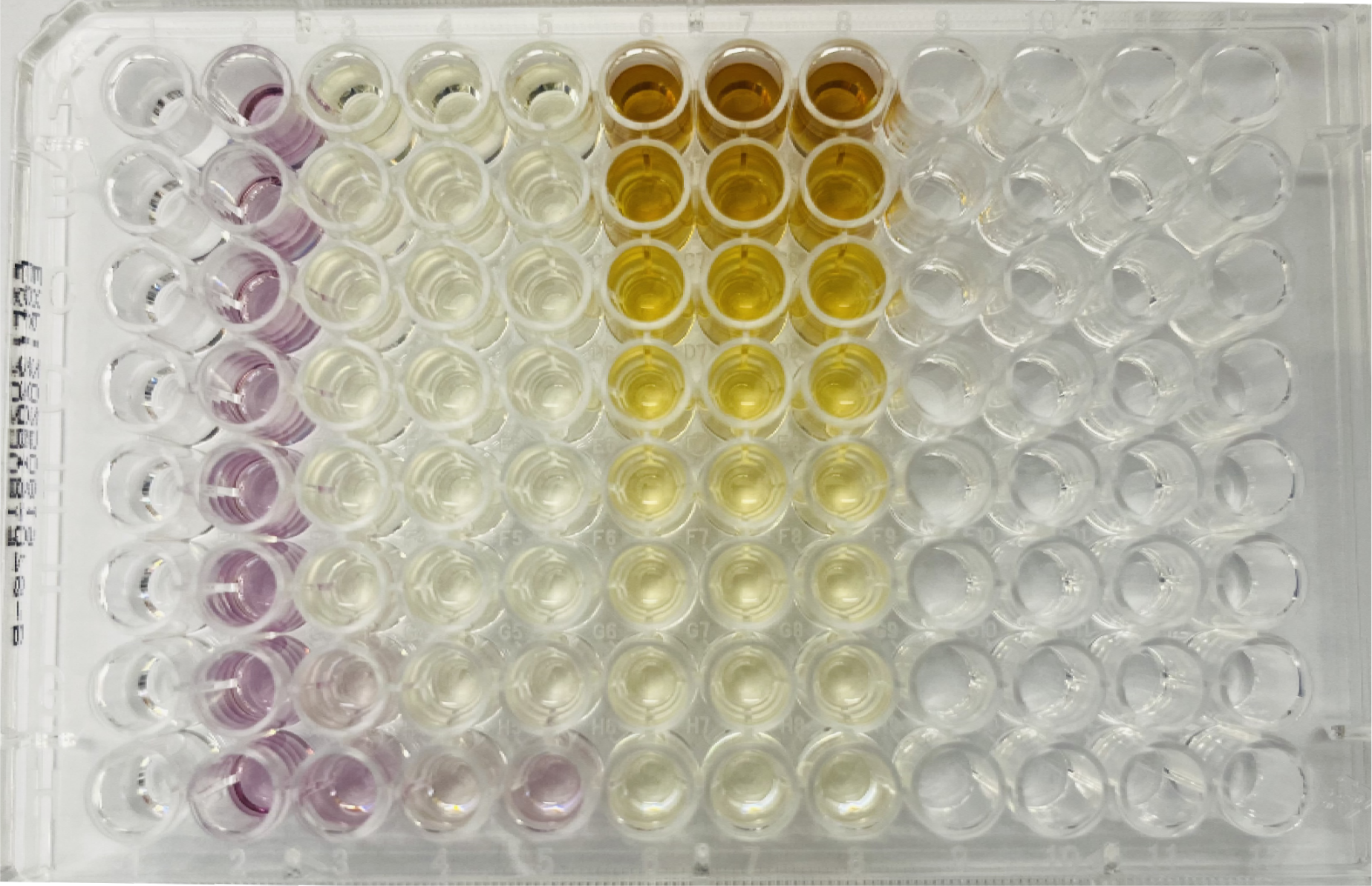
DPPH Assay C.

### Part III: Identify bioactive compounds/interactions via Mass Spec. & HPLC

The absorbance data was analyzed to compare the relative absorbance between the samples, the positive control, and DPPH. The statistical significance of populations was calculated to determine if differences in antioxidant potency were statistically significant. A one-way ANOVA with Tukey’s post hoc test was used. Then, we took the two most potent antioxidant samples and subjected them to LC-MS/MS analysis. This was done on a ThermoScientific LTQ XL mass spectrometer at the University of Rhode Island College of Pharmacy. We followed the procedures outlined in Jouaneh et al. 2022.^10^ Following analysis, we transferred raw mass spec data to mzXML files and followed a specialty pipeline developed by the Bertin laboratory at URI to analyze compounds in the antioxidant-rich extracts using the software platform available at gnps.ucsd.edu.^11^

### Additional considerations

Research required work with hazardous chemicals. Ethanol: flammable, potential fumes capable of eye irritation; 2,2-diphenyl-1-picrylhydrazyl (DPPH): can stain and damage skin, potential fumes capable of eye irritation.

To combat such risk, PPE was used: safety glasses, lab coats, and nitrile gloves. Work was done under a fume hood to mitigate the propagation of harmful fumes capable of irritating eyes, skin, or lungs. Furthermore, chemical waste was disposed of in waste containers in the supervisor’s lab. Safe practices were reviewed on chemical SDS^12,13^ for ethanol and DPPH. URI Environmental Health and Safety Protocols were also followed.^14^

## RESULTS

Following extraction and evaporation via inert nitrogen, the dried extract weights were determined. The dried *Camellia sinensis* extract had a mass of 45.5mg; *Averrhoa carambola* 34.6mg; *Salvia rosmarinus* 11.2mg; *Laurus nobilis* 8.2mg; and *Psidium guajava* 34.7mg (Figure 1). The dried extracts were then reconstituted to concentrations of 10mg/mL by adding appropriate amounts of water based on their respective powdered extract mass (Figure 1). Extracts were then compared in triplicate in decreasing concentrations.

With the addition of the DPPH solution to the sample triplicates, absorbance changes were observed in all compounds and compared to the positive control, quercetin, and the negative standard, DPPH. In a DPPH assay, a lesser absorbance indicates greater antioxidant activity. Initially, net absorbance was calculated (Figures 4.a, 4.b, 4.c, 7.a) and was then used to calculate individual factors of changes in each compound (Figures 5.a, 5.b, 5.c, 7.b) to normalize and prevent bias that may arise due to initial extract colors. *Factor of Change = Net / Initial.* For example, *Salvia rosmarinus* began and ended with a particularly dark extract color (Figure 3.b) that needed to be considered for an accurate comparison. Finally, the factor of changes were standardized as a relative percentage of the DPPH absorbance factor of change to best grasp relative absorbances (Figures 5.a, 5.b, 5.c, 7.c) and to best compare to the positive and negative controls. *Relative Factor of Change = Extract Factor of Change / DPPH Factor of Change*

**Figure 4.a.**
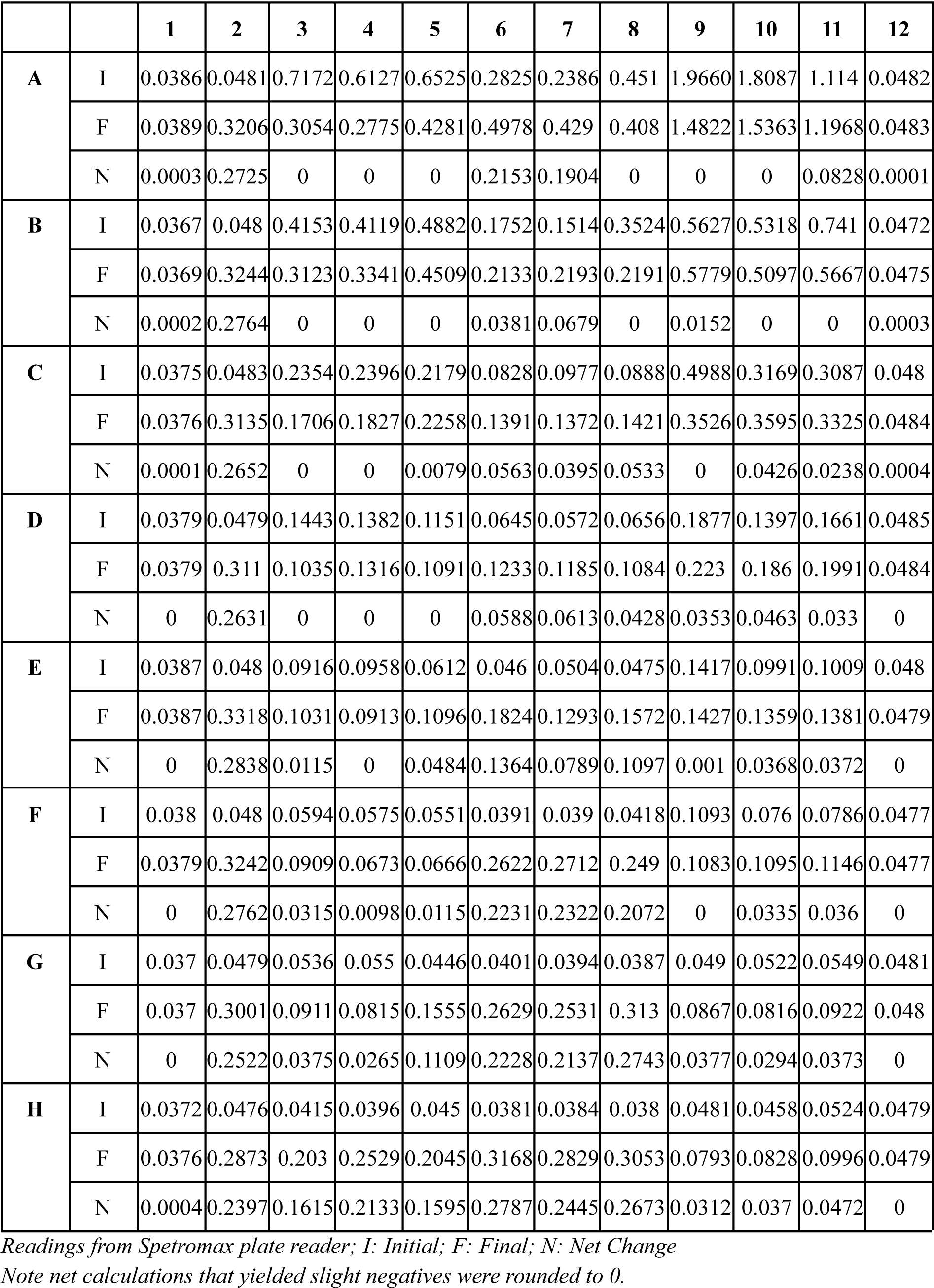
Raw Absorbance Data Plate Map A.

**Figure 4.b.**
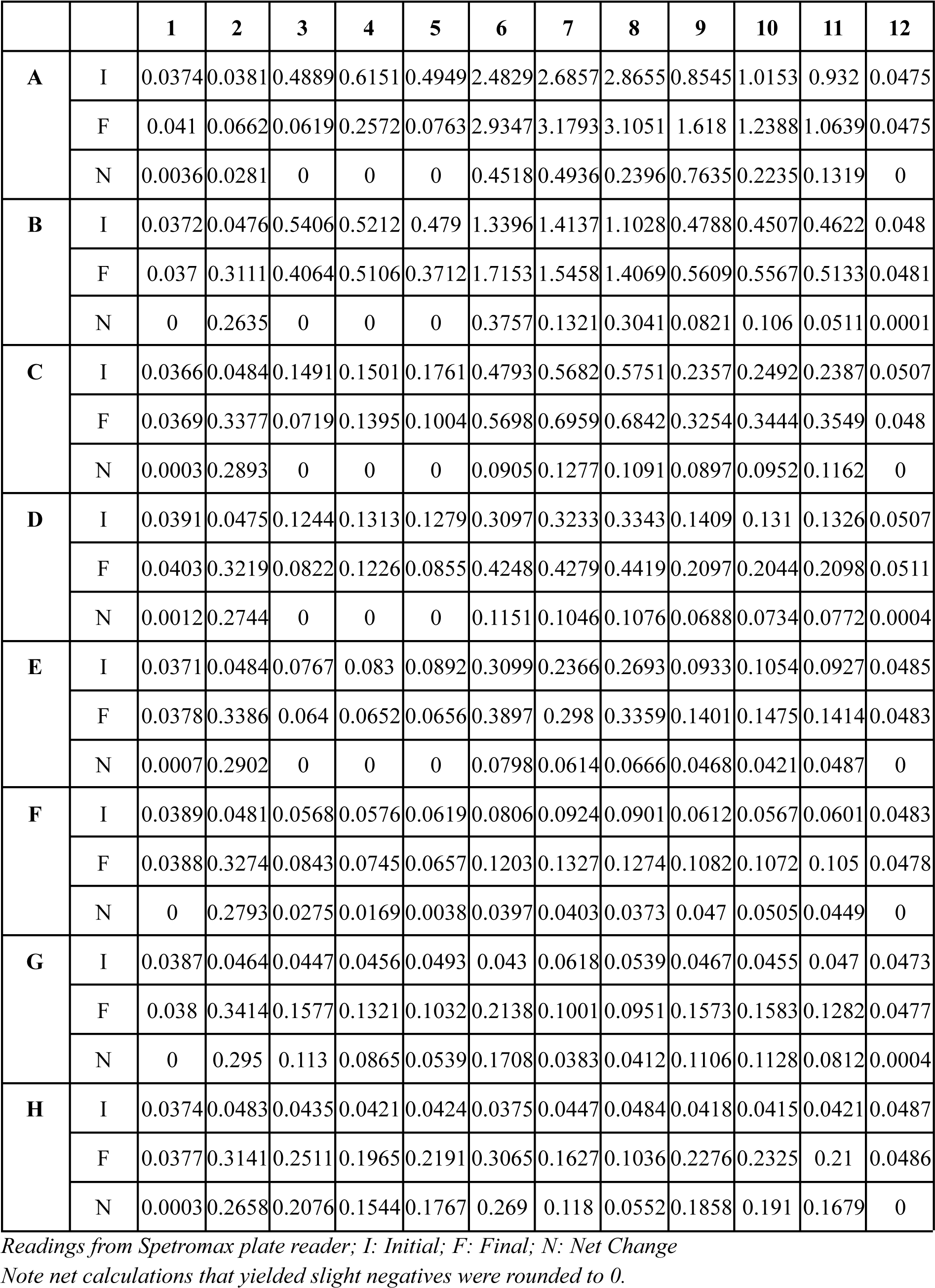
Raw Absorbance Data Plate Map B.

**Figure 4.c.**
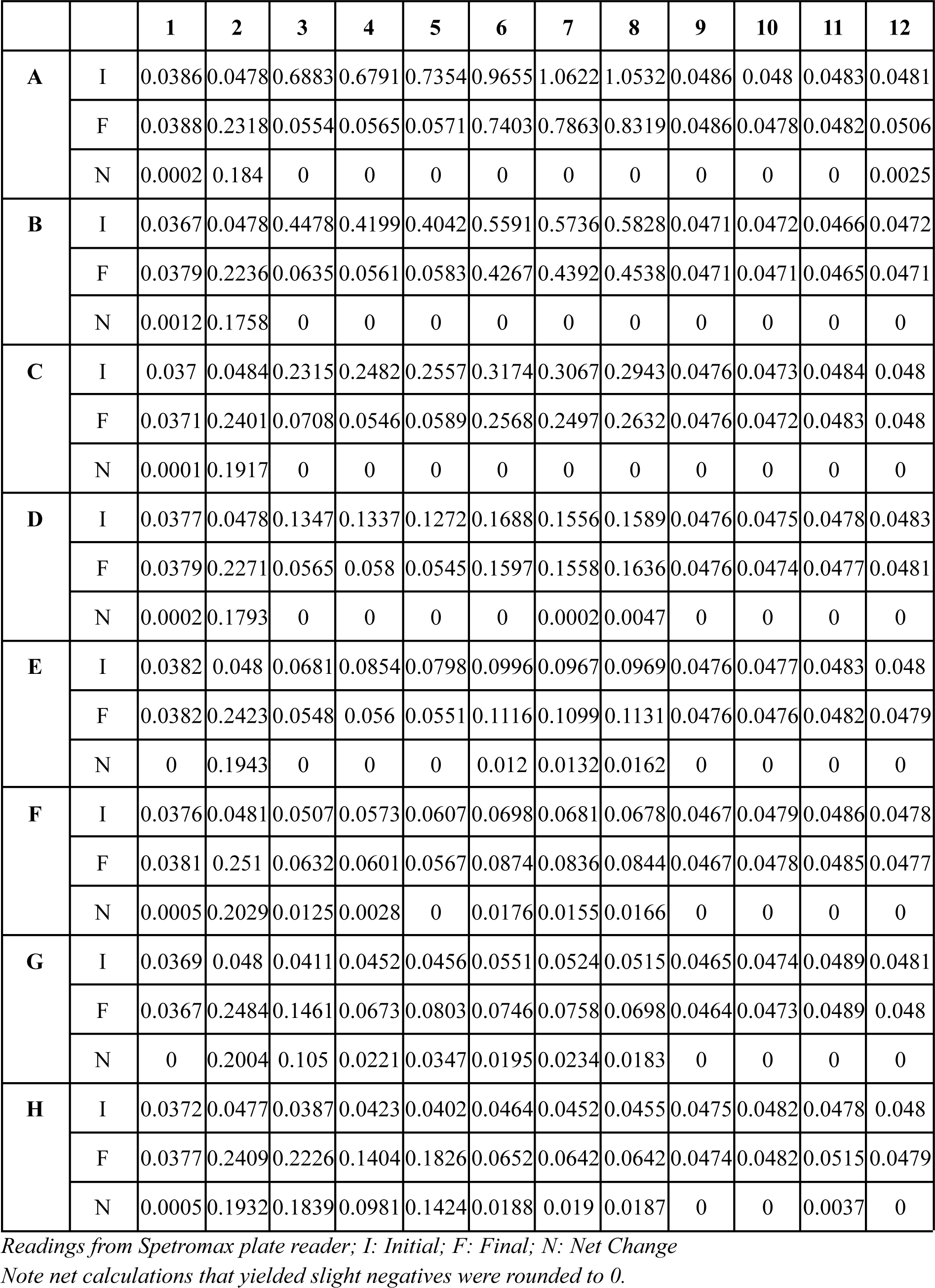
Raw Absorbance Data Plate Map C.

**Figure 5.a.**
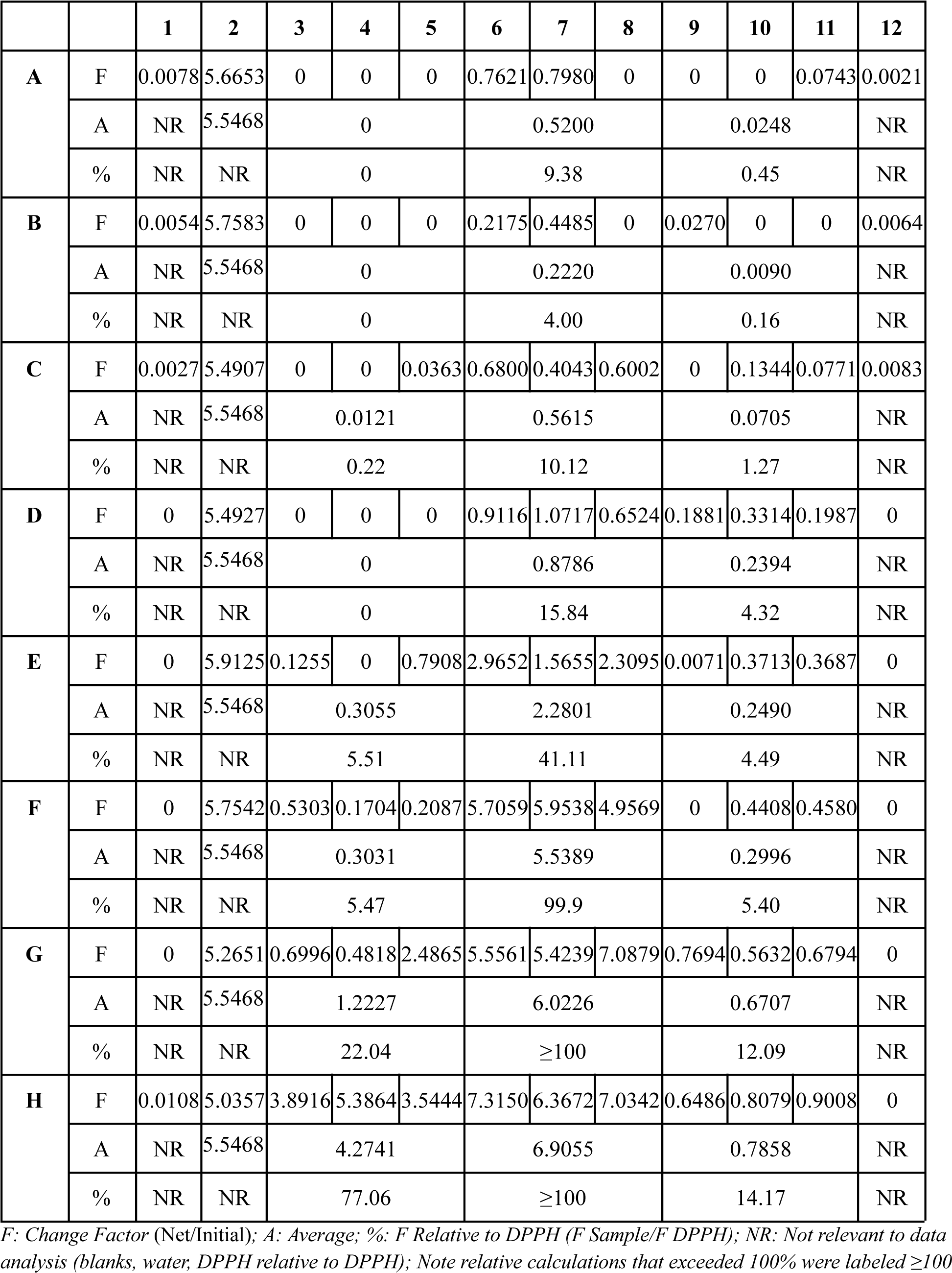
Absorbance Comparative Analysis Map A.

**Figure 5.b.**
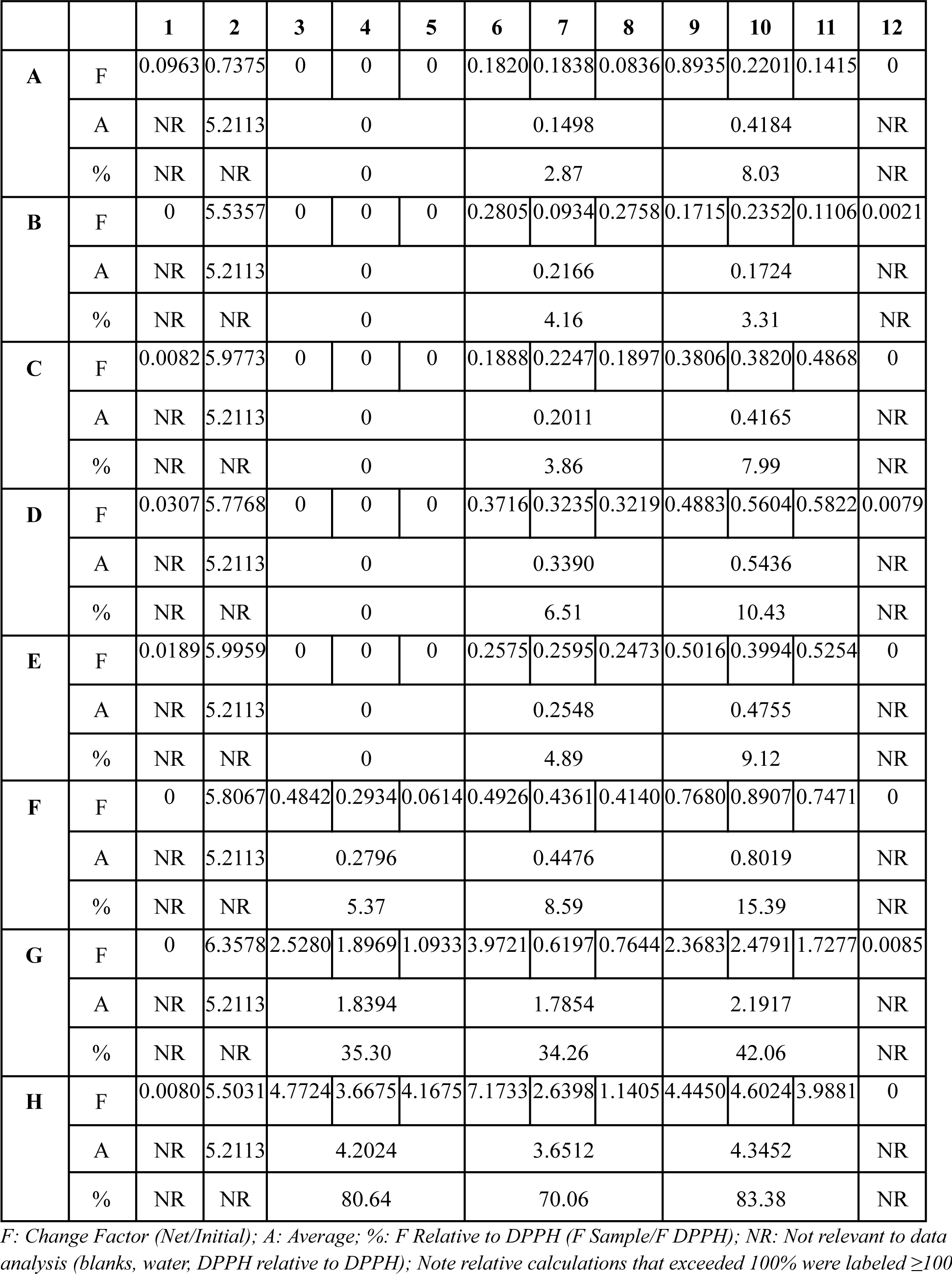
Absorbance Comparative Analysis Map B.

**Figure 5.c.**
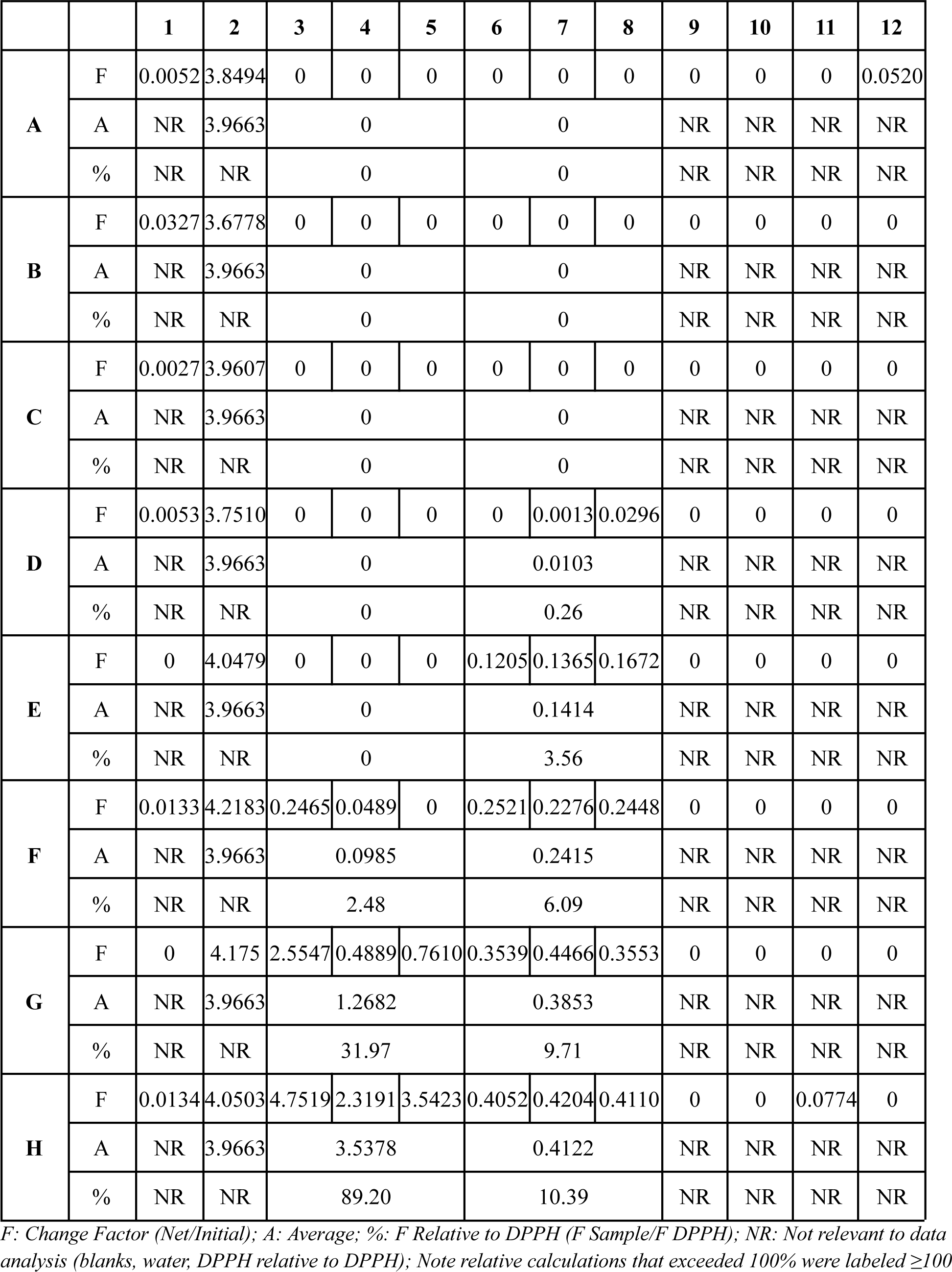
Absorbance Comparative Analysis Map C.

**Figure 6.a.**
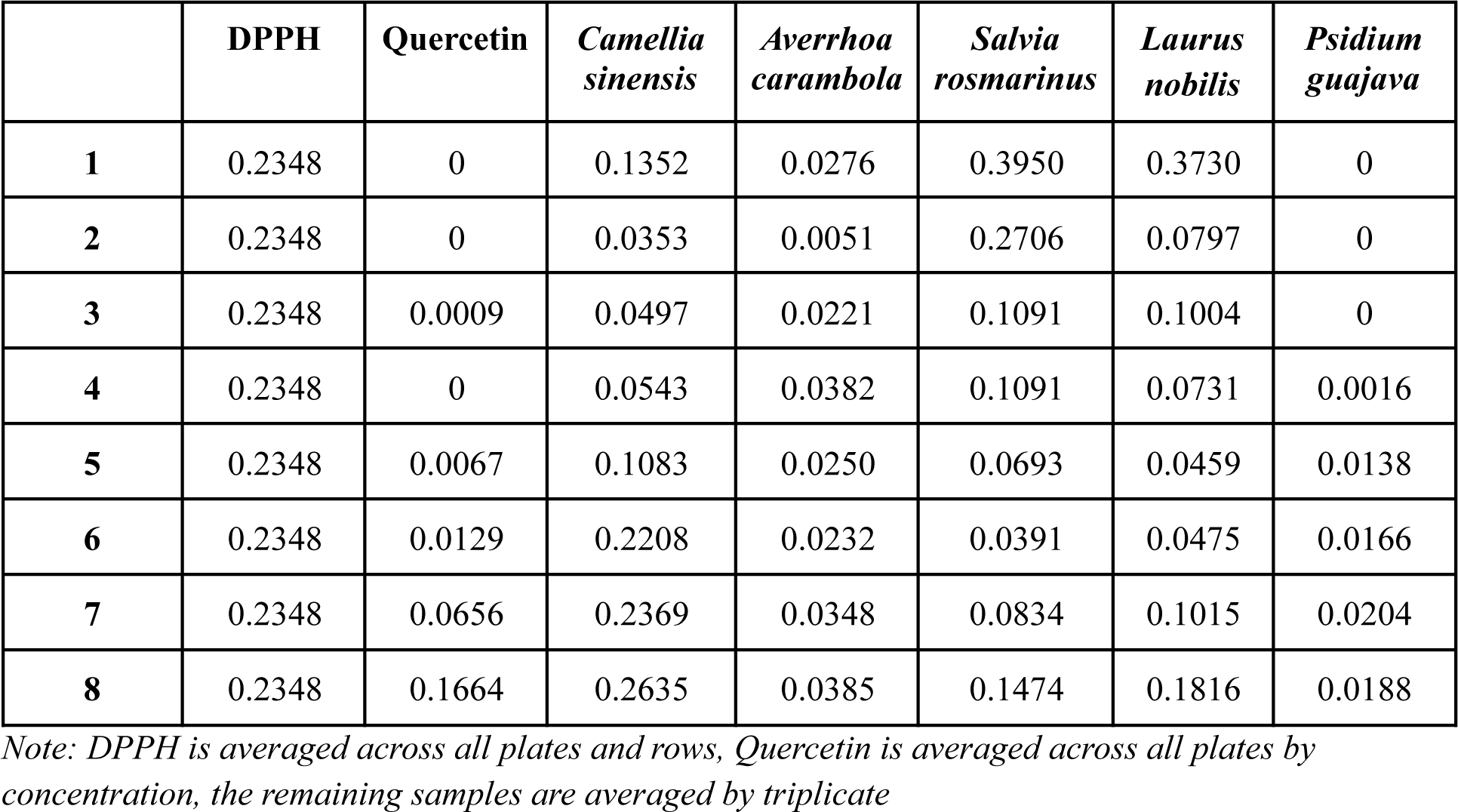
Combined Net Absorbance.

**Figure 6.b.**
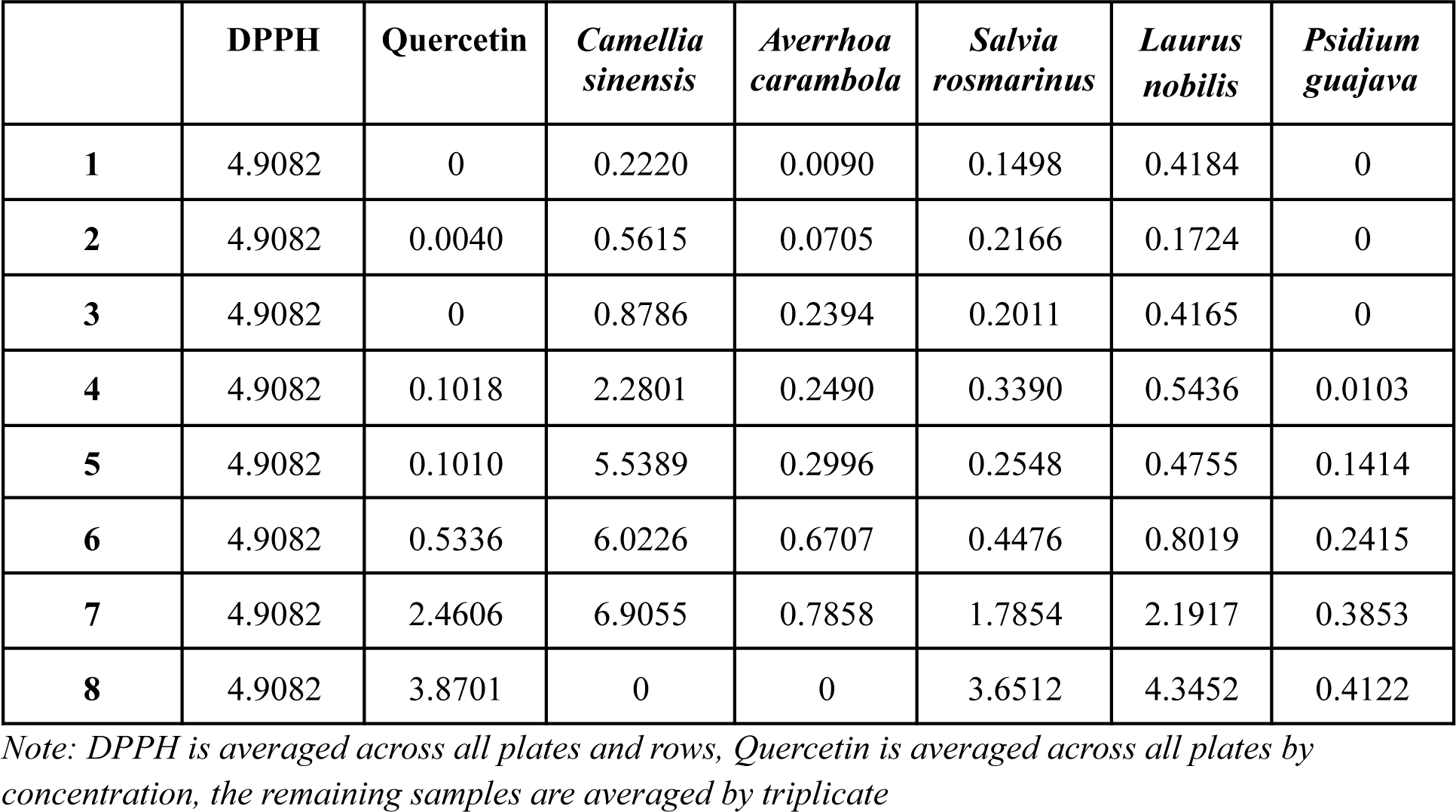
Combined Change Factor Absorbance.

**Figure 6.c.**
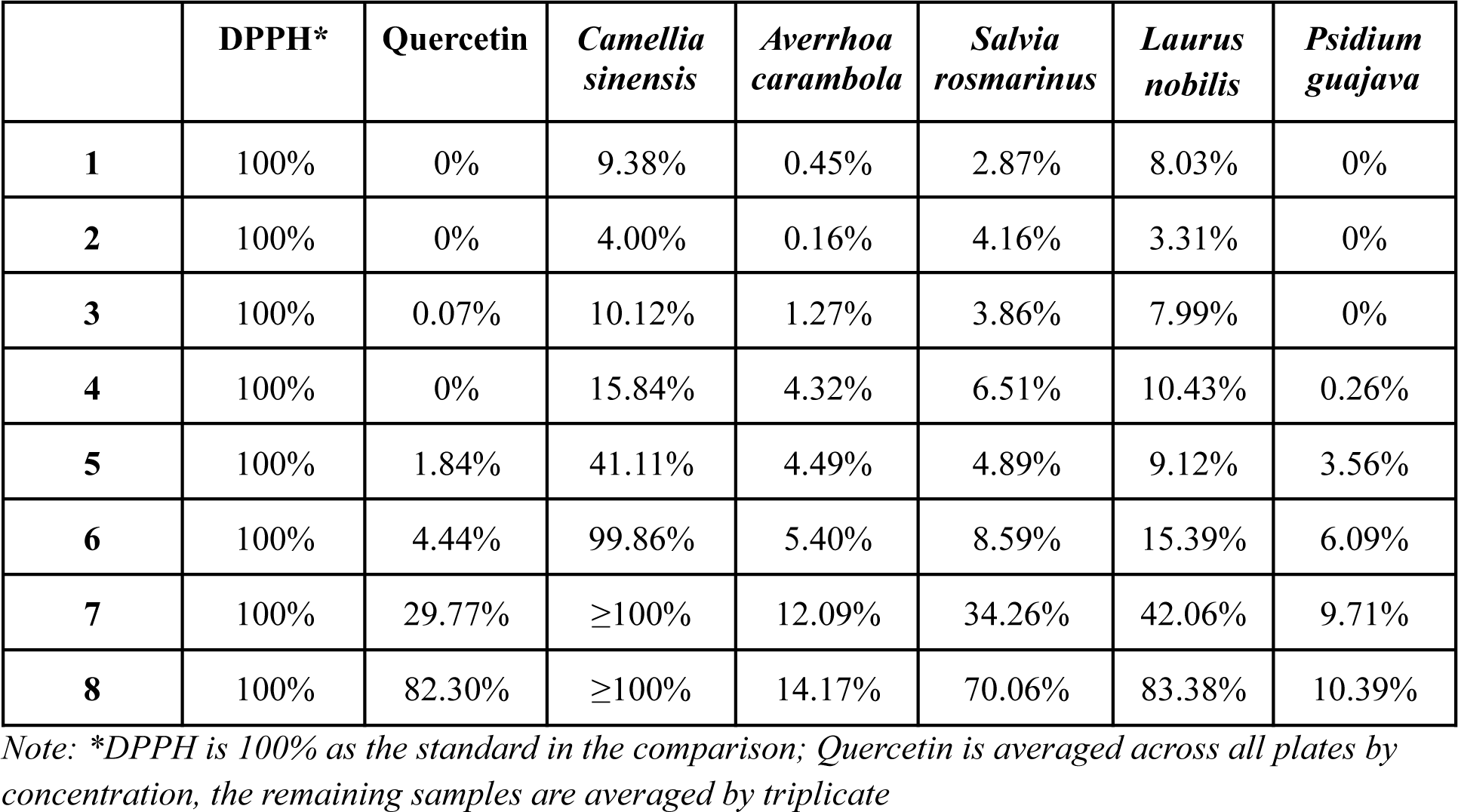
Combined Relative Absorbance.

**Figure 7.a.**
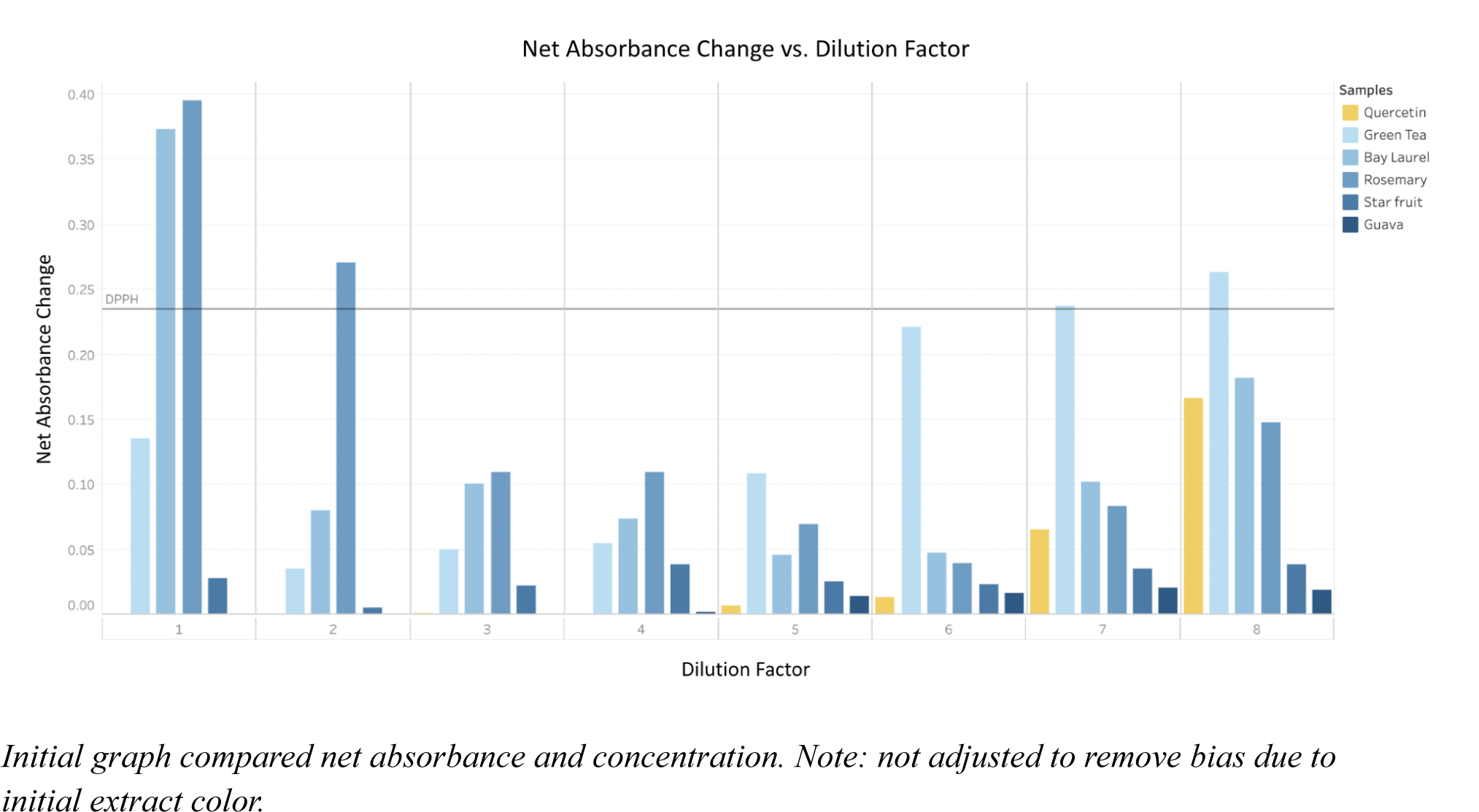
Net Absorbance vs Concentration Graph.

**Figure 7.b.**
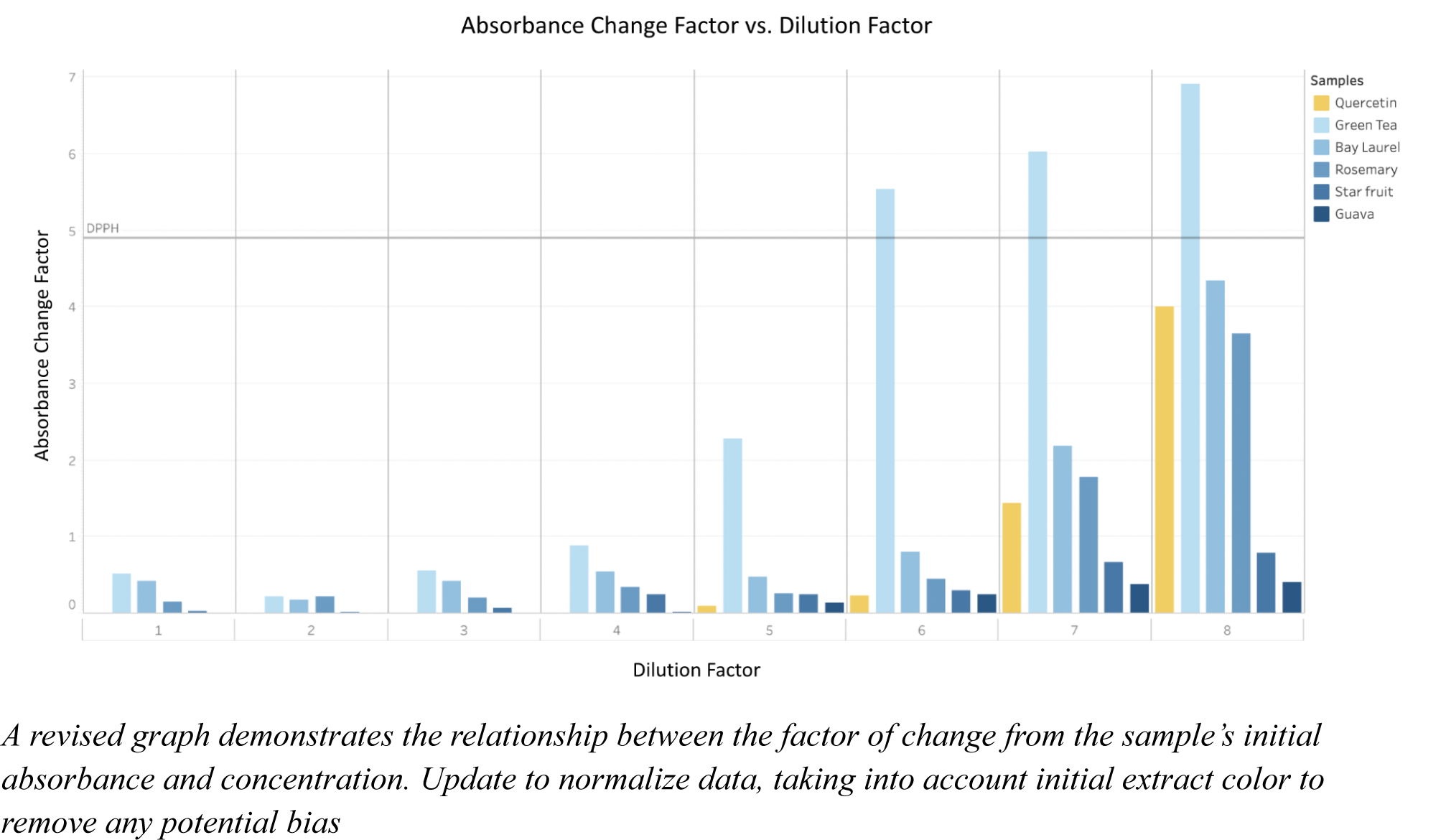
Change Factor Absorbance vs Concentration Graph.

**Figure 7.c.**
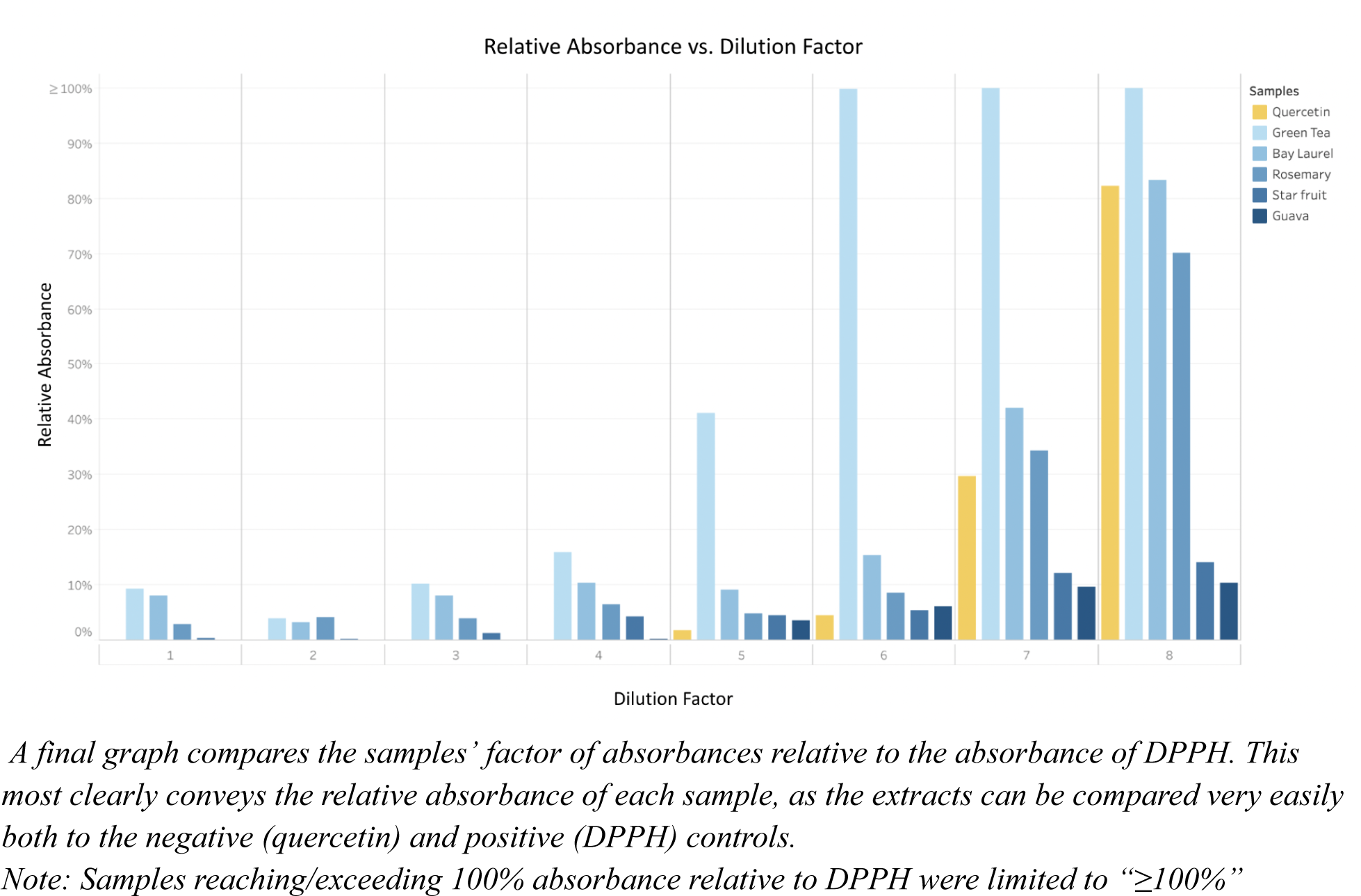
Relative (to DPPH) Absorbance vs. Concentration Graph.

At the first six dilution factor concentrations, all samples represented absorbance changes less than 100% of the DPPH standard (Figures 6.c, 7.c). Beyond six dilution factor concentrations, *Camellia sinensis* reached and/or exceeded 100% absorbance relative to DPPH, representing the greatest relative absorbance factor change out of the plant extract samples. *Laurus nobilis* followed, representing the second highest absorbance factor change out of all dilution factors, with an absorbance factor change 83.38% that of DPPH (Figures 6.c. 7.c) at the lowest extract concentration. Both *Camellia sinensis* and *Laurus nobilis* exceeded the absorbances of the positive control quercetin at all dilution factor concentrations (Figures 6.c. 7.c). The next greatest absorbance was *Salvia rosmarinus*, which also exceeded quercetin’s absorbance factor change at all dilution concentrations, excluding the greatest dilution factor when it measured 70.06% absorbance relative to DPPH (Figures 6.c, 7.c). However, this was not statistically significant (p > 0.05) per large error in the final dilution factor (Figures 8.a). Finally, representing the least degree of absorbance were the *Averrhoa carambola* and *Psidium guajava* samples, which at the greatest dilution factor possessed relative absorbances of 14.17% (Figures 6.c, 7.c) and 10.39% (Figures 6.c, 7.c), respectively, compared to the negative control DPPH. Both samples demonstrated a statistically significant (p < 0.05) difference in absorbance being substantially less than that of the positive control, quercetin, with a relative absorbance of 82.30% (Figure 6.c, 7.c). *Averrhoa carambola* and *Psidium guajava*’s relative absorbances were lower than that of all other samples and the positive control. This result was statistically significant (p < 0.05), though their difference between each other was not (Figure 8.a)

**Figure 8.a.**
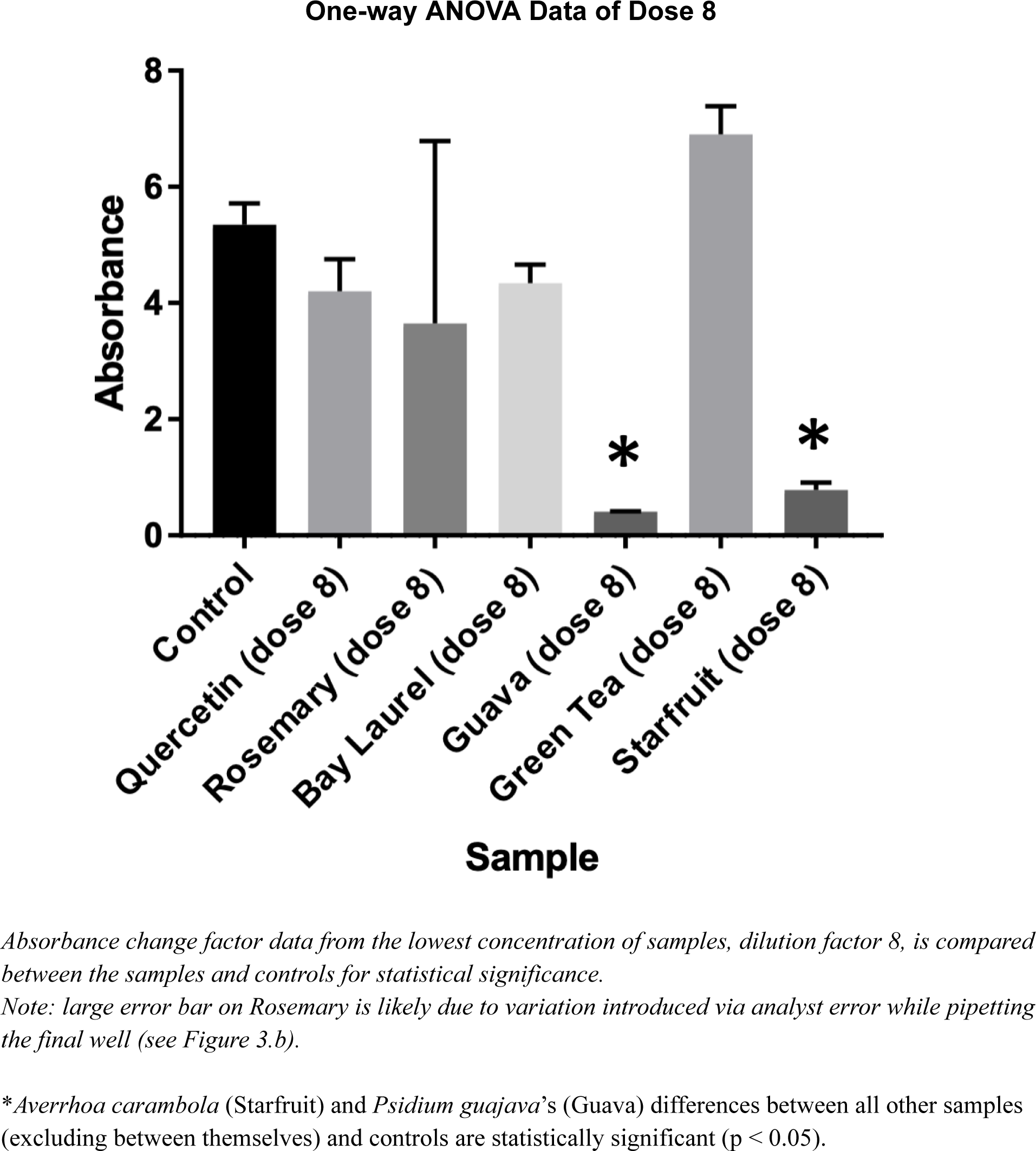
Statistical Significance Graph of Dose 8 (Dilution Factor 8)

Subsequently, the samples representing the least absorbance change upon the addition of DPPH, i.e., the samples representing the most potency in regards to antioxidant activity: *Averrhoa carambola* and *Psidium guajava*, were tested via LC-MS/MS analysis to identify potentially bioactive compounds. When compared to mass spectrometry libraries available at gnps.ucsd.edu, *Averrhoa carambola* was identified to contain vitexin-2’’-O-rhamnoside, isovitexin, saponarin, and a flavonoid glycoside: 8-[4,5-dihydroxy-6 -(hydroxymethyl)-3-(3,4,5-trihydroxy-6-methyloxan-2-yl)oxyoxan-2-yl]-7-hydroxy-2-(4-hydroxyphenyl) -5-[3,4,5-trihydroxy-6-(hydroxymethyl)oxan-2-yl]oxychromen-4-one (Figures 9.a, 10.a). Additionally, *Psidium guajava*’s mass spectra indicated the presence of quercetin and kaempferol (Figures 9.b, 10.b). These LC-MS compound identifications were distinct from that of the blank control, indicating the extract compound identifications distinctly align with their respective plant extracts (Figure 10.c).

**Figure 9.a.**
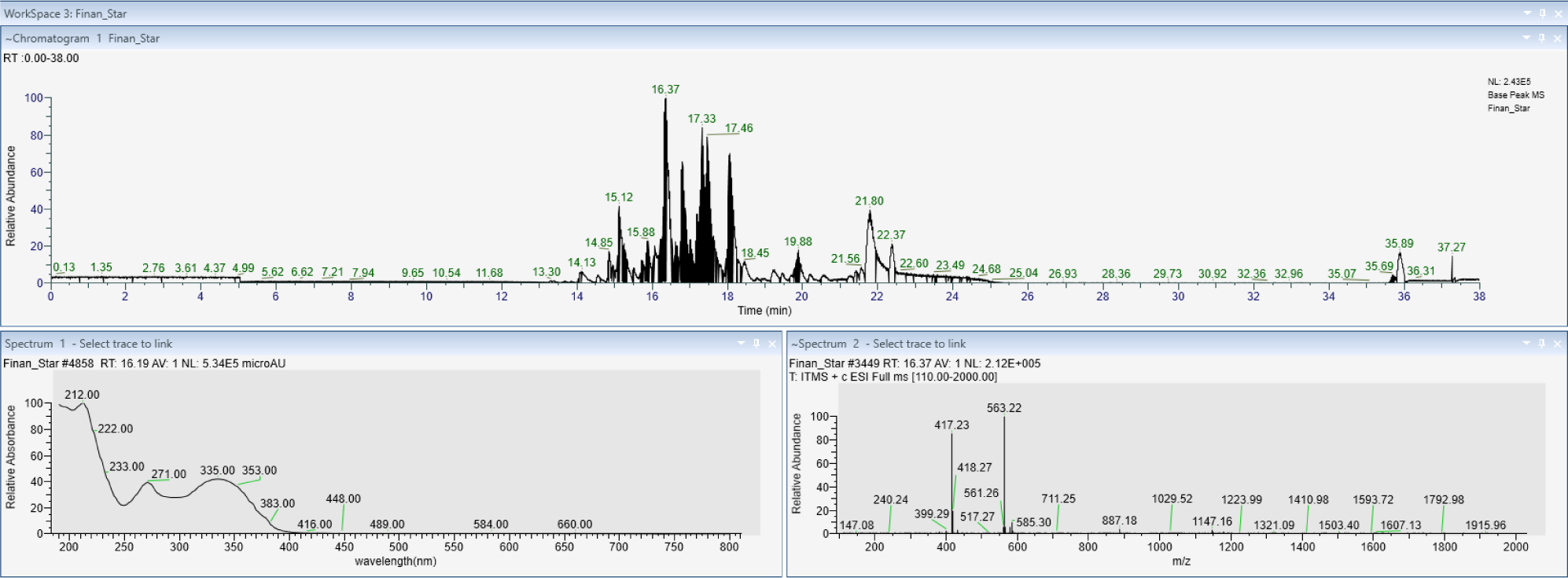
*Averrhoa carambola* Mass Spectra & HPLC.

**Figure 9.b.**
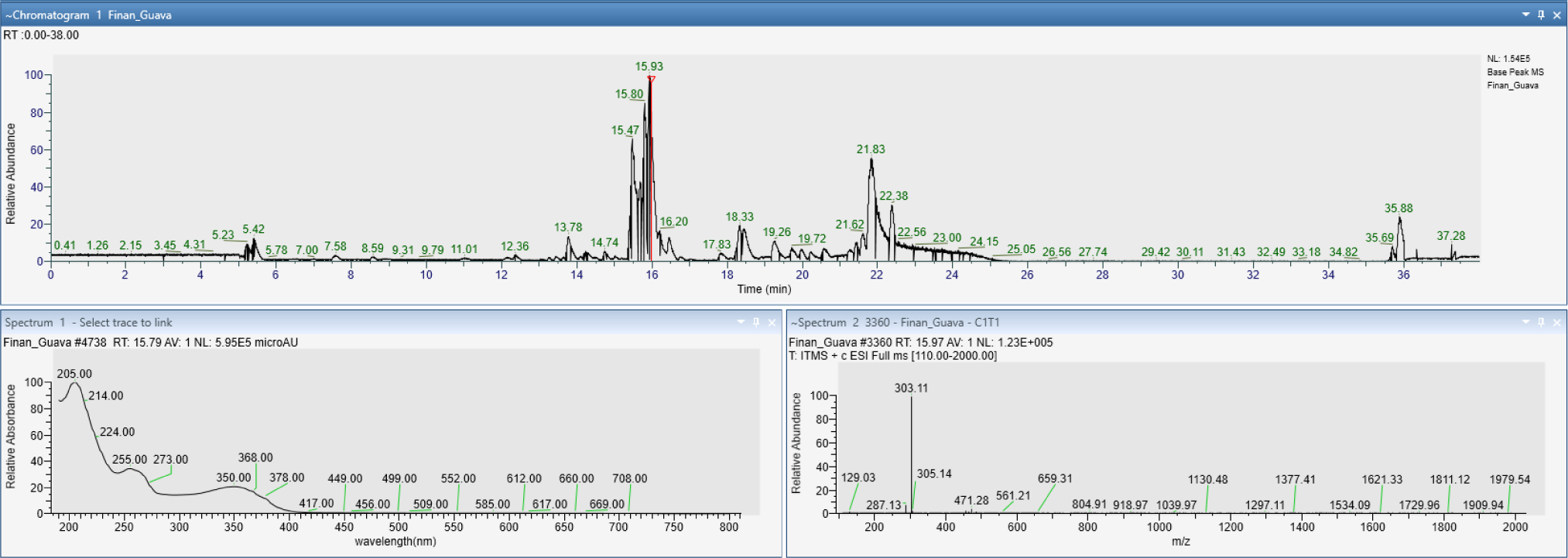
*Psidium guajava* Mass Spectra & HPLC.

**Figure 10.a.**
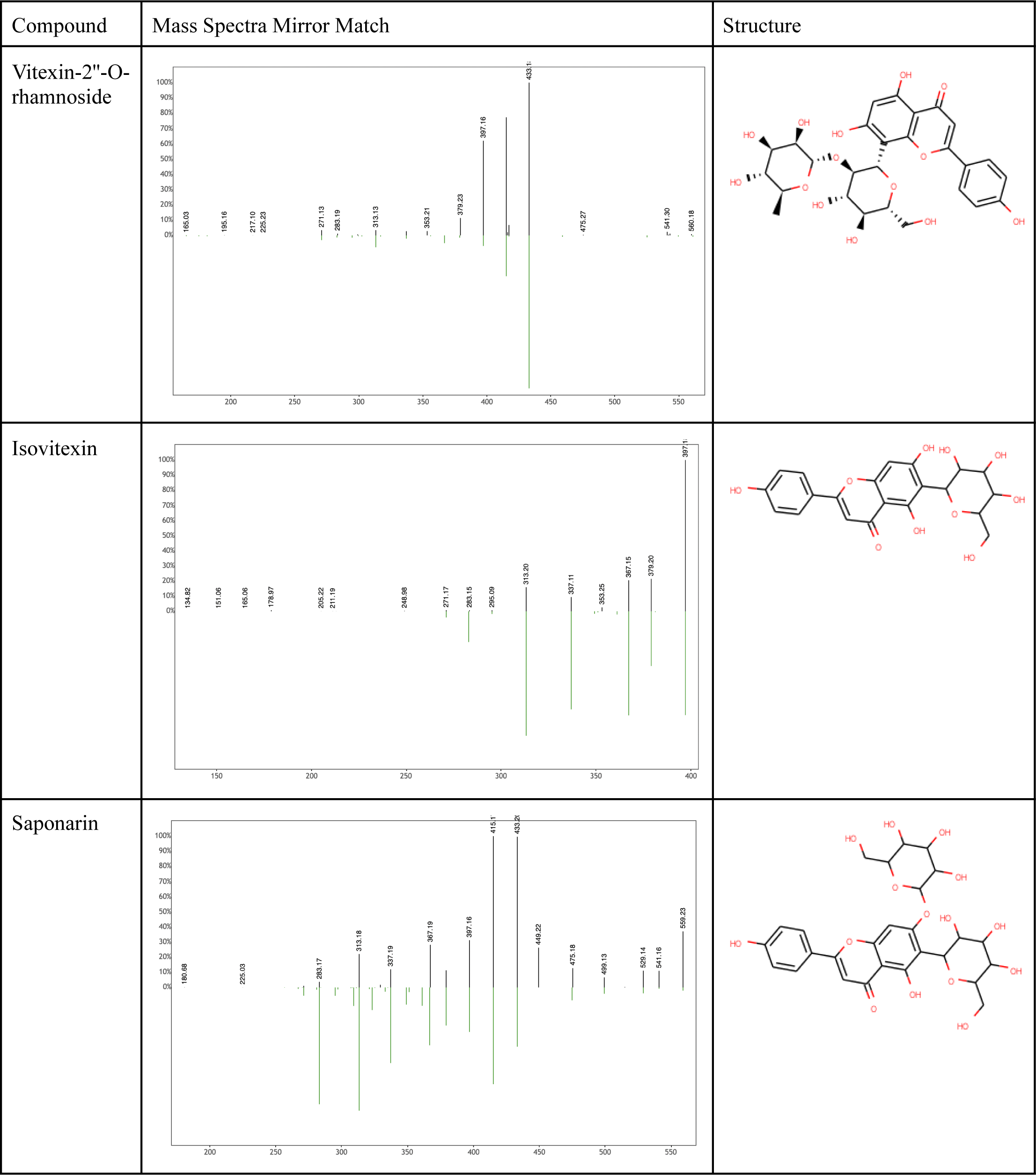
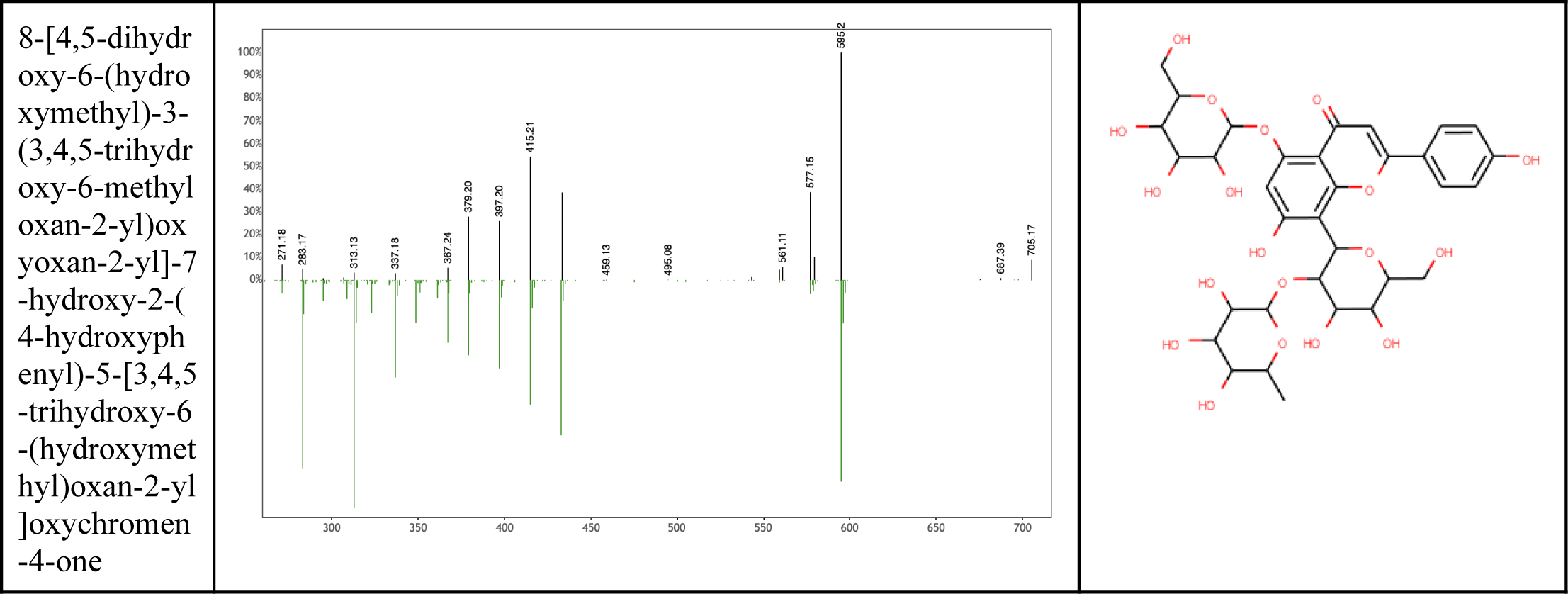
*Averrhoa carambola* Mass Spectra Analysis.

**Figure 10.b.**
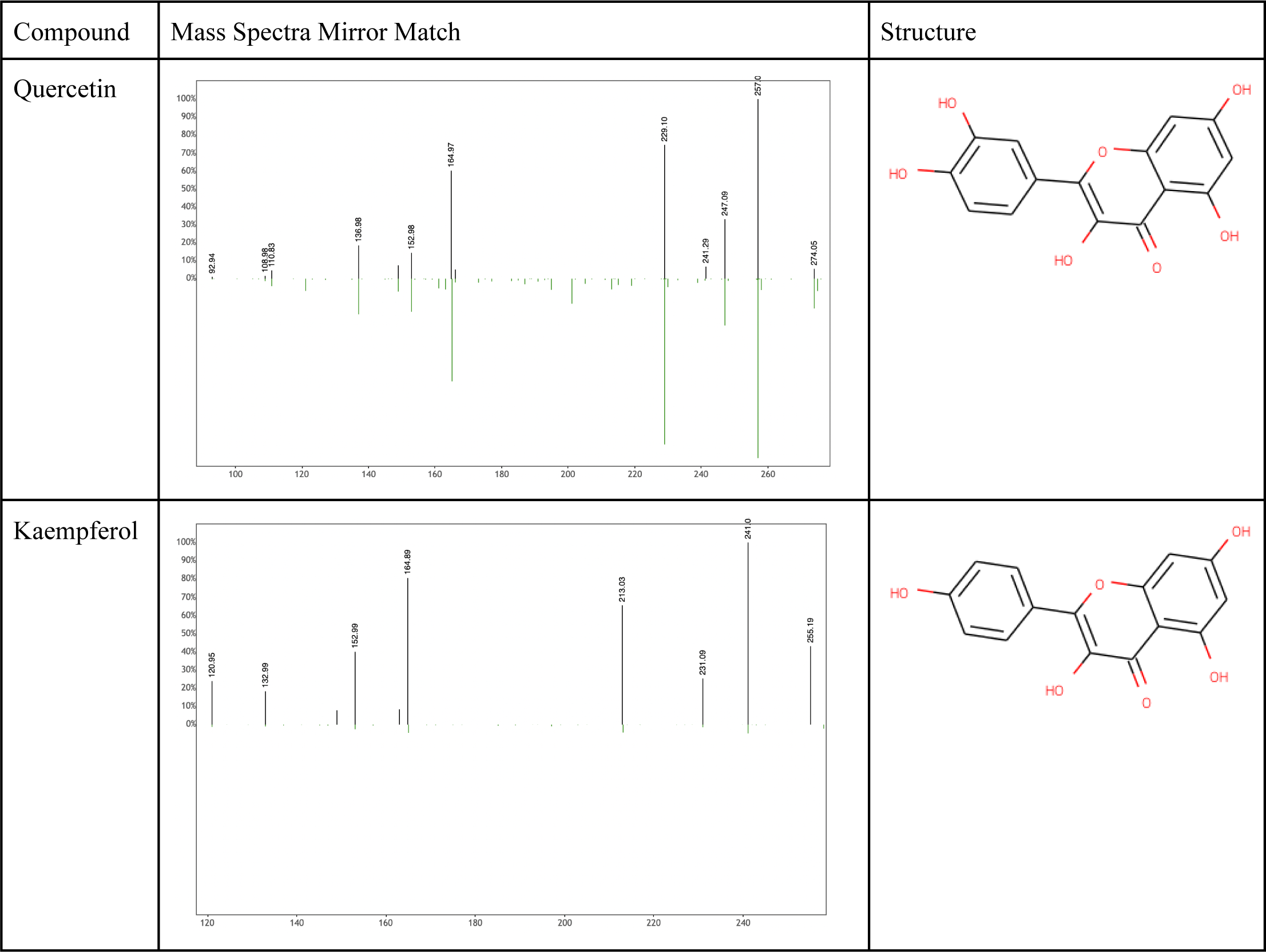
*Psidium guajava* Mass Spectra Analysis.

**Figure 10.c.**
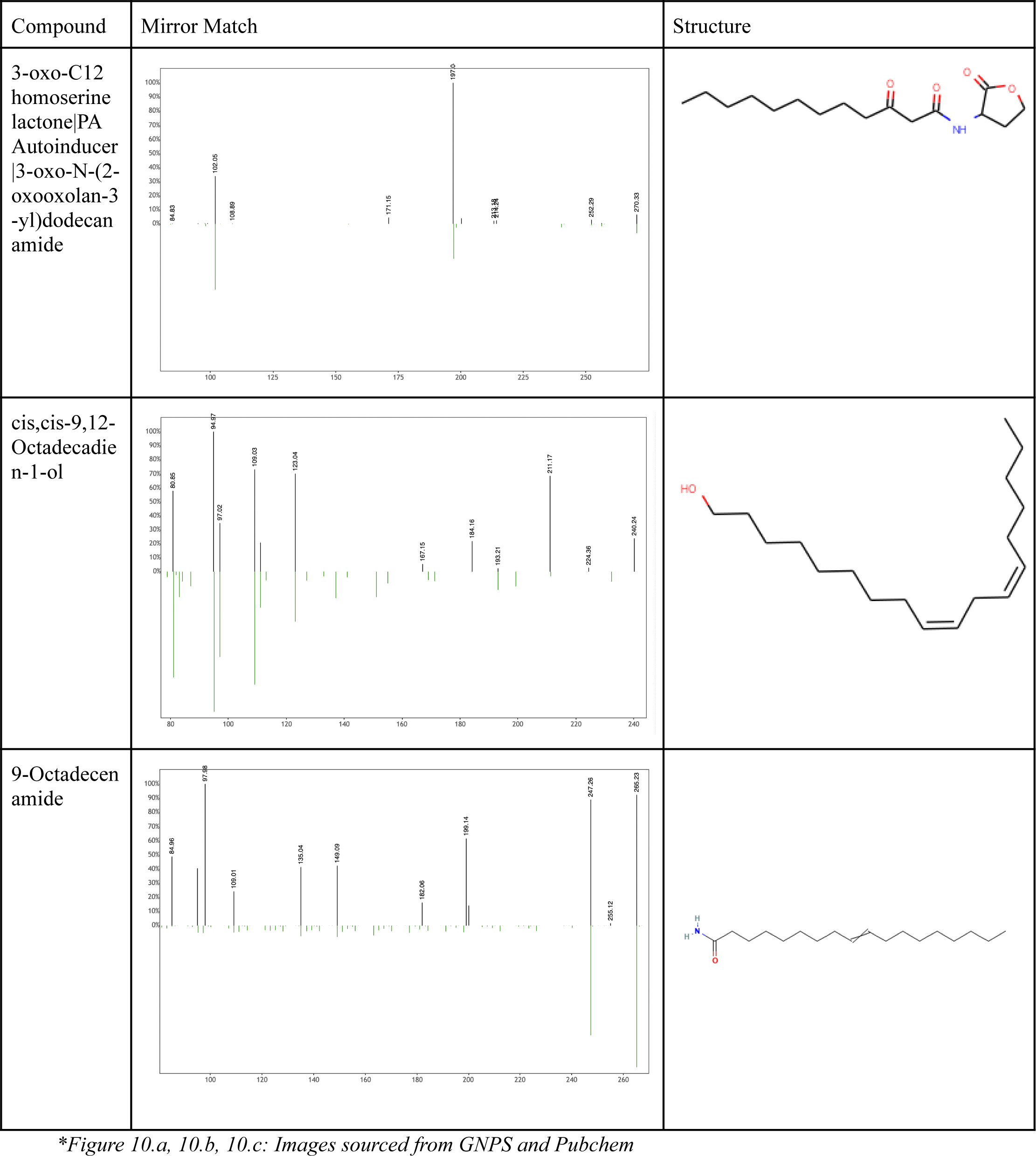
Blank Control Mass Spectra Analysis.

## DISCUSSION AND CONCLUSIONS

This study measured the relative potency of medicinal plant extracts in terms of antioxidant activity when compared to the positive control, quercetin, through absorbance changes in a DPPH assay. Representing the least degree of absorbance, hence the highest antioxidant potential, were the *Averrhoa carambola* and *Psidium guajava* samples, which at the greatest dilution factor possessed relative absorbances of 14.17% and 10.39% – less than that of quercetin, with a relative absorbance of 82.30% (Figure 6.c, 7.c). This would indicate greater antioxidant activity in these extract samples than even the positive control, as a lesser relative absorbance change would mean a greater neutralization of the purple-colored DPPH solution, which “is reduced to a yellow colored product, diphenylpicryl hydrazine.”^15^ Such neutralization of DPPH occurs through hydrogen atom transfer (HAT)^16,17^ but free radical scavenging can also occur through single electron transfer (SET), and chelation of transition metals.^17^ In DPPH, a hydrogen atom from the antioxidant species is transferred to DPPH’s nitrogen atom possessing an unpaired electron.^16^ Thus, our hypothesis of the antioxidant effects of the plant compounds is supported through the free radical scavenging witnessed in the DPPH assay.

Additionally, this ability of free radical scavenging contributes to the identified compounds’ properties and potential pharmacological applications. For example, kaempferol in *Psidium Guajava*, “augments the antioxidant potential of normal cells via modulating heme oxygenase (HO)-1 expression and mitogen-activated protein kinase (MAPK) pathways.”^18^ As a result, kaempferol offers numerous benefits as demonstrated through in vitro studies. This includes cancer-targeting properties in regard to apoptosis, angiogenesis, metastasis, and inflammation. Kaempferol has also been shown a remarkable affinity for malignant cells as opposed to healthy cells, reducing its risks of side effects.^19^ Furthermore, vitexin and its isomer isovitexin were identified in *Averrhoa carambola* and could potentially offer a variety of therapeutic applications in regard to their anticancer, antidiabetic, neuroprotective, cardioprotective, among other properties.^20^ Moreover, saponarin in *Averrhoa carambola* offers applications for diabetes through its ability to “restore liver function and architecture”^21^ and has also demonstrated relevance in sleep disorder treatments.^22^

However, the effectiveness of the aforementioned compounds is often substantially limited in regard to their bioavailability or lack thereof. Nanoparticle technology offers potential solutions to this issue, already demonstrating potential value in nanochemoprevention in the flavonoid Epigallocatechin -3-gallate (EGCG) which also offers potential anticancer properties.^19^ Additional research into the bioavailability of the aforementioned compounds should be conducted to determine their potential utility in modern medicine. Additionally, studies of the pharmacological effects of plant extract compounds are often limited to in vitro experiments and/or non-human tissue. Further experimentation should be conducted to determine the effectiveness of such compounds on human tissues and can only be fully substantiated through the lens of modern medicine through double-blind clinical trials. Corroboration of the claims of such bioactive molecules can standardize traditional herbal medicines prevalent in countries including China and India, ensuring safe and effective medical practices are upheld.

In summary, demonstrated efficacy in treatments of the potent antioxidant compounds in *Averrhoa carambola, Psidium guajava,* and beyond, can potentially yield revolutionary new treatments in a wide variety of diseases and medical practices.

